# Environmental specialization and cryptic genetic divergence in two massive coral species from the Florida Keys Reef Tract

**DOI:** 10.1101/2020.11.17.387522

**Authors:** John P. Rippe, Groves Dixon, Zachary L. Fuller, Yi Liao, Mikhail Matz

## Abstract

Broadcast-spawning coral species have wide geographic ranges, spanning strong environmental gradients, but it is unclear how much spatially varying selection these gradients actually impose. Strong divergent selection might present a considerable barrier for demographic exchange between disparate reef habitats. We investigated whether the cross-shelf gradient (nearshore - offshore - deep) is associated with spatially varying selection in two common coral species, *Montastraea cavernosa* and *Siderastrea siderea*, in the Florida Keys. Toward this end, we generated a *de novo* genome assembly for *M. cavernosa* and used 2bRAD to genotype 20 juveniles and 20 adults of both species from each of the three reef zones to identify signatures of selection occurring within a single generation. Unexpectedly, each species was found to be composed of four genetically distinct lineages, with gene flow between them still ongoing but highly reduced in 13.0-54.7% of the genome. Each species includes two sympatric lineages that are only found in the deep (20 m) habitat, while the other lineages are found almost exclusively on the shallower reefs (3-10 m). The two “shallow” lineages of *M. cavernosa* are also specialized for either nearshore or offshore: comparison between adult and juvenile cohorts indicates that cross-shelf migrants are more than twice as likely to die before reaching adulthood than local recruits. *Siderastrea siderea* and *M. cavernosa* are among the most ecologically successful species on the degraded Florida Keys Reef Tract, and this work offers important insight on the genomic background of divergent selection and environmental specialization that may in part explain their resilience and broad environmental range.

## INTRODUCTION

Oceans continue to warm at a pace that many worry will threaten the sustainability of tropical coral reefs within the next 50 to 100 years (Hoegh-Guldberg et al., 2007). Major coral bleaching events are occurring with increasing frequency, compounding the loss of living coral cover year after year (Hughes et al., 2017). The severity of this decline is prompting reef managers to consider proactive measures designed to enhance coral adaptation to heat stress. Examples of such measures include assisted gene flow, or the injection of adaptive alleles into threatened populations via hybridization with corals from warmer environments, and selective breeding and outplanting of individuals that demonstrate elevated resilience to thermal stress (Hagedorn et al., 2018; Quigley et al., 2019; van Oppen et al., 2015, 2017).

To predict the success of these actions and understand the evolutionary potential of coral reefs more generally, the patterns by which genetic variation is naturally distributed and exchanged among individuals on the reef must be evaluated. Coral populations on modern reefs have endured the selective pressure of decades of environmental change and therefore offer valuable insight into the factors driving the segregation and sorting of genetic diversity. Yet, while conventional surveys of broad-scale population genetics provide a useful introduction to the genetic landscape of a study species, often the most pertinent follow-up questions are left unresolved. What are the genetic processes underlying local adaptation and population differentiation in scleractinian (*i.e*., stony) corals? What mechanisms may be restricting the exchange of genetic material between divergent populations? To ensure lasting success in restoration activities and the efficient use of limited resources, it will be imperative to take these factors into account.

Complicating this task immensely is the incomplete and often muddled picture of speciation and diversification in corals. Rapid advances in DNA sequencing have ushered in a wave of genomic datasets, which have revealed that species boundaries originally defined based on morphological characteristics are often not consistent with inferences based on experimental and population genetic evidence (Willis et al., 1997). Interspecific hybridization, for example, is prevalent within many coral genera (reviewed in Willis et al., 2006). This has fueled debate on the relevance of reticulate evolution (Vollmer & Palumbi, 2002), or the notion that speciation in scleractinian corals is a fluid process involving backcrossing and migration among diverged lineages. On the other hand, apparent lack of gene flow and reproductive isolation within certain species has led to suggestions that cryptic speciation may be more common in corals and related taxa than is currently recognized (Bongaerts et al., 2010a; Ladner & Palumbi, 2012; Richards et al., 2016; Rosser, 2015; Schmidt-Roach et al., 2013; Warner et al., 2015). Together, these dynamics can have important implications on the exchange of genetic variation, and of adaptive alleles more specifically, among coral populations.

One important process that can lead to genetic divergence is local adaptation due to spatially varying selection across an environmental gradient. It was long presumed that due to large dispersal distances and few apparent physical barriers to gene flow, most marine organisms were well-mixed genetically across their range, with high migration rates largely swamping out the development of local adaptation (Ronce & Kirkpatrick, 2001). While there is evidence for such high gene flow in some marine systems, for many other species population connectivity is considerably lower than previously thought, suggesting that high gene flow is not a general trend (reviewed in Palumbi, 2004). In reef-building corals, local adaptation is often inferred from *ex situ* or reciprocal transplant experiments in which individuals tend to perform better in conditions most similar to their environment of origin (Barkley et al., 2017; Howells et al., 2013; Kenkel et al., 2013, 2015). Several recent studies have linked these patterns to a genetic basis (Dixon et al., 2015; Palumbi et al., 2014), but the specifics of this association are not well resolved. Thus, there is still much left unknown about the prevalence of local adaptation in stony corals and the underlying genetic mechanisms that maintain it.

Here we study two coral species, *Montastraea cavernosa* and *Siderastrea siderea*, which are ubiquitous in the wider Caribbean. We sampled two age cohorts (juveniles and adults) of each species across three sites representing a cross-shelf gradient in the lower Florida Keys: a shallow nearshore, shallow offshore, and deep offshore reef environment. Each reef zone represents a unique set of limiting environmental conditions that have a demonstrated effect on coral fitness (Kenkel et al., 2013; Lirman & Fong, 2007). With this in mind, we expected to see evidence of spatially varying selection among reef zones, manifesting as an increase in genetic differentiation in the adult cohort compared to the juvenile cohort. In particular, considering that both coral species are broadcast-spawners with the dispersal potential vastly exceeding the spatial scope of our sampling, we expected to see this effect only at a minority of loci in the genome involved in local adaptation.

## MATERIALS & METHODS

### Sample collection and site features

Samples were collected from two massive coral species, *Montastraea cavernosa* and *Siderastrea siderea*, from three sites along the cross-shelf gradient in the lower Florida Keys (Figure 1A). Each sampling site typifies a unique environmental setting: (1) a shallow nearshore patch reef (3-5 m) adjacent to Summerland Key on the shoreward boundary of Hawk Channel, (2) a shallow offshore site (~10 m) representing a backreef community adjacent to the relict barrier reef structure of the Florida Keys Reef Tract, and (3) a deep offshore site (~20 m) near the base of the reef slope seaward of the Looe Key Sanctuary Preservation Area. These habitats differ substantially in their ambient environmental conditions, in that nearshore reefs experience on average greater terrestrial influence (*i.e*., higher turbidity, dissolved nutrient levels and chlorophyll a) and larger temperature fluctuations than offshore reefs (Lirman & Fong, 2007). Exposure to the Florida Current maintains a relatively stable temperature regime in both offshore habitats, although greater depth is associated with lower levels of photosynthetically active radiation (Lesser, 2000).

**Figure 1.**
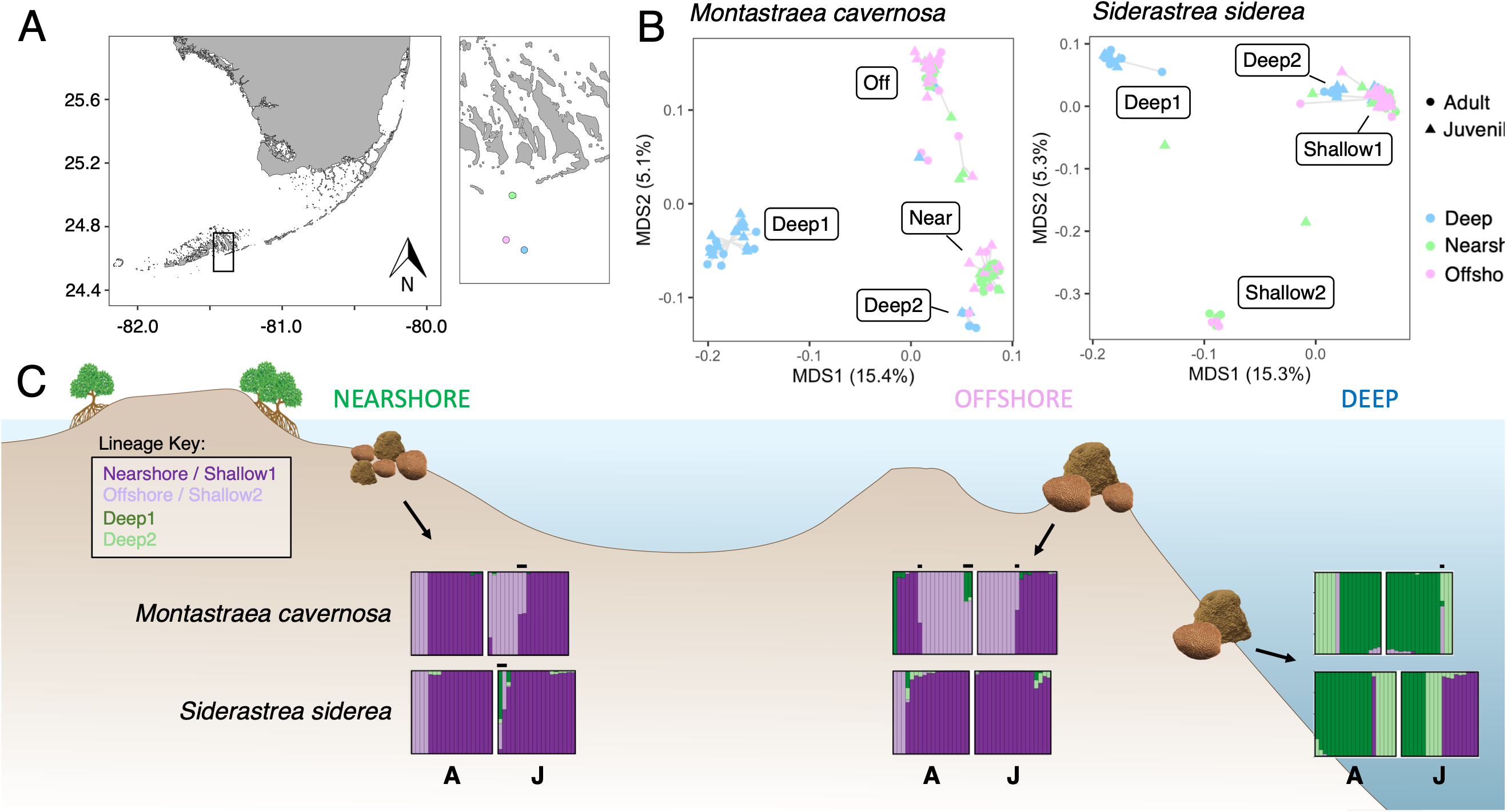
Sample map and spatial population genetics. (**A**) The three sampling sites (Nearshore: green, Offshore: pink, Deep: blue) are displayed in the inset map. (**B**) Principal coordinate analyses (PCoA) demonstrate the four distinct lineages evident within each species. The colors of individual points correspond to the habitats specified in the sample map. (**C**) Admixture plots for each species are displayed along the bathymetric cross-section diagram, separated by adults (A) and juveniles (J) in each habitat. Note again the four distinct lineages in each species, two distributed primarily across the shallow nearshore and offshore sites and two restricted to the deep. Throughout the text, lineages are referenced as follows: Nearshore / Shallow1 (dark purple, *M. cavernosa / S. siderea*), Offshore / Shallow2 (light purple, *M. cavernosa / S. siderea*), Deepl (dark green), Deep2 (light green). Hybrids with >25% ancestry to a secondary lineage are indicated with black bars above the plots.

At each sampling site, small tissue fragments were collected from 20 adults and 20 juveniles of each species when possible. In some cases, 20 individuals of both life stages could not be located, leaving a total sample size of 118 for *M. cavernosa* and 123 for *S. siderea*. Life stages were distinguished based on colony diameter, with colonies larger than 30 cm across classified as adults, and those smaller than 3 cm classified as juveniles. All samples were preserved in 100% ethanol at −20°C until transport to the University of Texas at Austin, where they were kept at −80°C until processing.

### Montastraea cavernosa *genome assembly and annotation*

One specimen of *M. cavernosa* was collected for genome sequencing from the West Bank of the Flower Garden Banks Marine Sanctuary (27.88° N, 93.83° W). We constructed a *de novo* genome assembly of *M. cavernosa* using a combination of PacBio reads and Illumina paired-end reads with 10X Genomics Chromium barcodes. The PacBio sequencing (9 SMRT cells, yielding 1.9 million subreads with N50 = 8kb, 15.7 Gb total) was performed at the Duke Center for Genomic and Computational Biology. 10X barcode libraries were generated and paired-end reads sequenced on an Illumina HiSeqX platform at the New York Genome Center.

Our assembly approach follows a similar method used to generate a reference genome for the coral *Acropora millepora* (Fuller et al., 2020). Briefly, we first constructed an initial draft assembly using only PacBio long reads with canu v1.6 (Koren et al., 2017). We then performed two rounds of scaffolding on this PacBio-only assembly using the 10X paired-end reads with scaff10x v1.1 (https://github.com/wtsi-hpag/Scaff10X). The resulting assembly underwent additional scaffolding using the 10X barcodes and paired-end reads with the ARCS/LINKS pipeline (Warren et al., 2015; Yeo et al., 2018). The PacBio long reads were then mapped to the assembly and used to join contigs with SSPACE-longread v1.1 (Boetzer & Pirovano, 2014). A final two rounds of scaffolding were then performed using scaff10x v1.1 again. We filled in gaps with PBJelly v15.8.24 (English et al., 2012) and error corrected with PacBio reads using the Arrow v5.0.1.9585 algorithm (https://github.com/PacificBiosciences/GenomicConsensus). To create the final assembly, we used the paired-end reads with 10X barcodes removed to polish with pilon v1.22 (Walker et al., 2014). In total, this resulted in 10,835 scaffolds with an N50 of 248Kb and maximum length of 1.8Mb.

An initial all-vs-all alignment of the assembly revealed the presence of redundant contigs. To further refine the assembly, we used the HaploMerger2 (Huang et al., 2017) pipeline to rebuild both haploid sub-assemblies from the potential polymorphic diploid genome assembly. Afterwards, the final phased 448Mb assembly contained 5,161 sequences, with an N50 of 343Kb and average length of 86kb. The maximum scaffold length was unchanged.

Protein coding gene models of the *M. cavernosa* genome were annotated using MAKER-P v2.31.9 (Campbell et al., 2014). MAKER-P was implemented using the pipeline available at the CyVerse website (http://www.cyverse.org/), available as an Atmosphere image and MPI-enabled for parallel processing. To predict gene models, we used previously published transcriptome data (SRA accession: SRP063463) (Kitchen et al., 2015) and mapped these reads using a reference-based assembly strategy with the TopHat v.2.1.1 and Cufflinks v2.2.1 pipeline (Trapnell et al., 2012), resulting in 43,577 transcripts. In addition, we generated a *de novo* transcriptome from the raw RNA-seq data with Trinity v2.0.2 (Grabherr et al., 2011), producing 200,233 transcripts. Both sets of transcripts were used as evidence in the MAKER-P pipeline.

For protein-level evidence in the annotation pipeline, we used peptide sequences from 8 robust coral species: *Pseudodiploria strigosa, Seriatopora hystrix, Platygyra carnosus, Montastraea faveolata*, *Madracis auretenra*, *Fungia scutaria*, *Favia sp*, and *Montastraea cavernosa* itself (Bhattacharya et al., 2016). We excluded other relatively evolutionary distant coral species due to the significant divergence between genomes, as they are likely to provide only limited evidence for gene annotation.

Before running MAKER-P, we also performed *de novo* repetitive element identification for the genome assembly using RepeatScout v1.0.5 (Price et al., 2005). The assembly was then soft masked with this resulting repeat library using RepeatMasker (http://www.repeatmasker.org). 44.7% of the genome was masked and identified as repetitive. To generate the final gene set, we performed three rounds of training and predicting gene models with MAKER-P, in combination with AUGUSTUS (Hoff & Stanke, 2019). In total, 30,390 non-redundant gene models were produced from the pipeline. We assessed the completeness and quality of the predicted final gene set using 978 near-universal single-copy orthologs from the Metazoan set available in BUSCO v3 (Simão et al., 2015; Waterhouse et al., 2018). Of these, 793 are completely present and 96 are present as fragments, for a total of 90% matches to BUSCO groups searched. Finally, functional annotation was performed for the gene set using the Trinotate pipeline (Bryant et al., 2017) and eggNOG-mapper (Huerta-Cepas et al., 2017).

### Reduced representation library preparation and sequencing

Genomic DNA was extracted from tissue samples using a modified phenol-chloroform procedure. First, samples were transferred to an extraction solution comprising 800 μL CTAB extraction buffer (2% CTAB, 100 mM Tris pH 8, 20 mM EDTA, 1.4 M NaCl), 1.6μL beta merceptoethanol, 1μL proteinase K, and 1μL RNAse A. Tissue was separated from the skeletal matrix and macerated by bead beating in a BioSpec MiniBeadbeater-96 for 15 seconds using 150-212 μm glass beads and then incubated at 42°C for at least one hour. Skeletal fragments were pelleted by spinning in a tabletop centrifuge at maximum speed for 15 minutes. The aqueous phase was then transferred to a clean tube, and 800 μL phenol-chloroform/isoamyl alcohol was added and vortexed into solution for 2-3 seconds. Samples were centrifuged at maximum speed at 4°C for 20 minutes to separate the DNA from organic contaminants. The aqueous supernatant was transferred to a clean tube to which 550 μL ice cold isopropanol was added and gently mixed by inverting. After incubating at −20°C for 20 minutes, samples were again centrifuged at 4°C and maximum speed for 20 minutes to pellet DNA. The supernatant was discarded, 1 mL 80% ethanol added, and the sample was centrifuged at 4°C and maximum speed for 5 minutes. The supernatant was discarded and sample tubes were allowed to air dry for 15-20 minutes. Lastly, pelleted DNA was dissolved in 30 μL warm (65°C) nuclease-free water.

Samples were then prepared for reduced-representation sequencing using the 2bRAD methodology (Wang et al., 2012). The latest version of the protocol, employing the triple-barcoding scheme and degenerate tags for identification of PCR duplicates, is available at https://github.com/z0on/2bRAD_denovo. Isolated DNA was purified using the Zymo Clean and Concentrator-10 kit and normalized to a concentration of 12.5 ng/μL. 50 ng DNA was then digested in a reaction comprising 1 U BcgI restriction enzyme, 1X NEB Buffer #3, 20 μM SAM in a total reaction volume of 6 μL. Digests were incubated at 37°C for one hour followed by 20 minutes at 65°C to inactivate the enzyme. Barcodes were then attached in two stages, first by ligation to the digested DNA fragments and secondly via PCR amplification. Ligation reactions consisted of 1X T4 ligase buffer, 400 U T4 ligase, and 0.25 μM of two double-stranded ligation adaptors, one of which contained an internal barcode, combined with 6 μL digested DNA in a total reaction volume of 20 μL. Ligation reactions were held at 4°C for 12 hours followed by 20 minutes at 65°C to inactivate the enzyme. Samples with unique ligation barcodes were then pooled and a second barcode was attached via PCR with thermocycler conditions set to 30 seconds at 70°C, followed by 15 cycles of 20 seconds at 95°C, 3 minutes at 65°C and 30 seconds at 72°C. PCR was carried out using 10 μL of pooled ligations with 1X Titanium Taq polymerase, 1X Titanium Taq buffer, 200 μM dNTPs, 0.12 μM unique ILL-BC barcode, 0.12 μM TruSeq-UN barcode, and 0.2 μM each of P5 and P7 adaptors in a total reaction volume of 50 μL. After confirming successful amplification, all samples were pooled and sequenced on the Illumina HiSeq 2500 platform at the University of Texas at Austin DNA Sequencing Facility.

### Data filtering and genotyping

Raw sequencing reads were trimmed of adaptors and demultiplexed using a custom script (https://github.com/z0on/2bRAD_denovo) and were quality filtered using the program cutadapt v1.14 (Martin, 2011), removing any reads with a Phred score of less than 15 at either end. Because *S. siderea* lacks a published genome assembly, a reference was generated *de novo* for genotyping. To do so, sample reads were first mapped to an aggregate genome comprising the four *Symbiodiniaceae* genera *Symbiodinium*, *Breviolum*, *Cladocopium* and *Durusdinium* (Aranda et al., 2016; Dougan, 2020; H. Liu et al., 2018; Shoguchi et al., 2013) using the program bowtie2 v2.3.4 (Langmead & Salzberg, 2012). Any reads that mapped successfully with a minimum end-to-end alignment score of −22.2 were removed so that those left behind could be assumed to belong to the coral host. The program cd-hit v4.8.1 (Fu et al., 2012) was then used in combination with custom Perl scripts (https://github.com/z0on/2bRAD_denovo) to cluster and assemble reads from the 60 samples with the highest remaining sequencing coverage into a reference genome with 30 equally-sized pseudo-chromosomes for mapping.

Using this *de novo* assembly for *S. siderea* and the *M. cavernosa* genome described above (https://matzlab.weebly.com/data--code.html), sample reads were mapped to the appropriate reference using bowtie2 (Langmead & Salzberg, 2012). Samples with greater than 25% of reads successfully aligned and at least 5X sequencing depth for more than 25% of loci were retained for further analyses. Those failing to pass both filters were removed (*S. siderea*: n=5, *M. cavernosa*: n=6). Alignment files were sorted and converted to BAM format using SAMtools v1.6 (Li et al., 2009).

All genotyping and identification of single nucleotide polymorphisms was performed using the ANGSD program suite (Kim et al., 2011; Nielsen et al., 2012; Korneliussen et al., 2014), which is well suited for low-coverage genetic data as it operates on genotype likelihoods, rather than hard genotype calls, in most downstream applications.

Lastly, prior to any population genetic analyses, samples representing genetically identical individuals (*i.e*., generated via asexual reproduction) were identified based on the degree of genetic dissimilarity as compared to technical replicates included in the sample set (*i.e*., separate sequencing libraries prepared from the same tissue sample). To do so, variant sites (biallelic SNPs) were identified under a stringent filtering scheme in order to ensure confidence in polymorphic loci—all loci were required to have a base call quality greater than Q25, mapping quality greater than Q20, a SNP *p*-value less than 1 x 10^-5^, a minimum minor allele frequency of 0.05 and be sequenced in at least 80% of samples. Based on these loci (*M. cavernosa*: 8,039 SNPs, *S. siderea*: 13,016 SNPs), pairwise identity-by-state (IBS) distance was calculated between all samples, and a hierarchical clustering tree was constructed using function *hclust* in R v3.6.1 (R Core Team, 2019) to distinguish genetically identical individuals. Any samples exhibiting a genetic dissimilarity as low as known technical replicates were assumed to be genetically identical, and only the individual of each clonal group with the greatest total read count was chosen to be retained for further analyses (n = 2 samples removed per species; Figure S1). All the scripts and detailed outline of the bioinformatic procedures are available at https://github.com/z0on/2bRAD_denovo.

### Population structure

Genome-wide population structure was assessed in each species using two methods: (1) individual admixture proportions among K inferred ancestral lineages, and (2) principal component analysis based on IBS distance. First, variant sites were identified using the *ANGSD* filters described above—here, the requirement for a locus to be sequenced in 80% of samples reflects the reduced sample size after genetically identical individuals were removed (*M. cavernosa:* 8,265 SNPs, *S. siderea*: 14,827 SNPs). Based on variation at these loci and given a specified number of ancestral lineages (K = 2-6) in the population, the proportion of each individual’s ancestry derived from each inferred lineage was estimated using the program ngsAdmix (Skotte et al., 2013). Additionally, to compare and corroborate the interpretation of this analysis, pairwise IBS distance between individuals was calculated in *ANGSD*. Principal coordinate analysis (PCoA) was performed on the resulting matrix using the *vegan::capscale* function in R (Oksanen et al., 2019) to visualize the clustering of samples with respect to the major axes of genetic variation. The distinct genetic lineages identified for each species based on the combination of these results form the basis for all subsequent analyses.

### Modeling demographic histories

To infer the demographic history of these genetic lineages, we used two approaches based on allele frequency spectra (AFS), both aiming to reconstruct demographic scenarios that most accurately reproduce the observed AFS. For both analyses, target loci were identified using modified filters in ANGSD in order to retain all well-sequenced sites, including invariant ones (Matz, 2018). Modifications include increasing the minimum sequencing and mapping quality to Q35 and Q30, respectively, while removing the minimum allele frequency and minimum SNP p-value filters. These additional filters returned roughly a million sites and did not qualitatively change our conclusions of population structure (Figure S2). AFS were then generated using ANGSD for the four genetic lineages that were identified in each species, using only samples that had less than 25% of alternative lineage ancestry as determined by ngsAdmix (*M. cavernosa*: n = 99 (7 putative hybrids removed), *S. siderea*: n = 110 (2 putative hybrids removed)).

Since the reference genome for *M. cavernosa* was sampled from an outlying population both in terms of geographical distance and habitat type (near-mesophotic deep habitat in the Flower Garden Banks in the Gulf of Mexico), we assumed the alleles not matching the reference to be derived in the populations studied here. To correct for possible misidentification of ancestral and derived allelic states, two-dimensional demographic models included the proportion of misidentified ancestral states as a free parameter. Since *S. siderea* lacks a reference genome, alleles cannot be polarized into ancestral and derived states; thus, all AFS were “folded” at a minor allele frequency of 0.5.

To reconstruct the history of effective population size changes for each genetic lineage, we applied StairwayPlot v2 (X. Liu & Fu, 2015) to one-dimensional AFS. StairwayPlot is an unsupervised analysis that does not require pre-specified demographic models. Prior to this analysis, we have applied BayeScan (Foll & Gaggiotti, 2008) to identify *F*_ST_ outliers (*i.e*., sites showing more allele frequency divergence between genetic lineages than expected under neutral island model). BayeScan was run using default settings: 20 pilot runs (5,000 iterations in length) followed by a burn-in of 50,000 iterations and a final run of 100,000 iterations. Sites that were assigned a q-value of <0.5 for being an *F*_ST_ outlier were removed, leading to removal of 6.4 % and 5.2% of all sites for *M. cavernosa* and *S. siderea*, respectively. Such permissive q-value threshold was chosen to ensure that all potentially non-neutral sites were removed, following Matz et al. (2018).

Additionally, we used the *Moments* Python library (Jouganous et al., 2017) based on twodimensional AFS to estimate not only population size changes, but also the timing and magnitude of introgression between each pair of genetic lineages. Unlike StairwayPlot, the Moments modeling workflow requires demographic models to be specified at the outset, which are defined and executed using custom Python scripts. Here, we specified a set of 108 different models that represent a wide range of possible demographic scenarios—notable differences between models include variations in the occurrence and timing of symmetric or asymmetric migration, the timing and dynamics of population size changes, and crucially, whether migration rates are allowed to vary across the genome to represent “islands of differentiation” (Duranton et al., 2018).

The *Moments* procedure followed three main steps to ensure confidence in model selection and parameter estimation. (1) First, ten bootstrapped two-dimensional AFS were generated using ANGSD for each pairwise population comparison. For *M. cavernosa*, bootstrapping was done by resampling genomic scaffolds; for *S. siderea* - pseudo-chromosomes from the read-based reference. For all ten AFS, each of the 108 demographic models was run six times with randomly chosen initial parameter values to ensure that each model converges to its optimum on each replicate at least once. (2) Following Burnham and Anderson (2003), model log likelihoods were then converted to Akaike’s Information Criterion (AIC) using the formula:

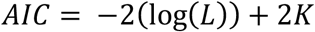

where *L* is the likelihood of the model and *K* is the number of model parameters. The best-fit run of each model (out of six) on each bootstrap replicate was identified, and the “winning” demographic model was identified as the model with the lowest median AIC across the ten bootstrap replicates. (3) Lastly, the winning model was fitted to 100 bootstrapped AFS in order to estimate the confidence of demographic parameters. Again, six random restarts per bootstrap replicate were performed, each of them initiated by randomly perturbing the parameters estimated at the model selection stage. All *Moments* models, accessory scripts, and instructions for this procedure are available at https://github.com/z0on/AFS-analysis-with-moments.

Estimated parameters (*T, θ, M*) were converted to time in years (*t*), effective population size in number of individuals (*N_e_*), and migration rates as the fraction of the total population that are new immigrants per generation (*m*) assuming a mutation rate of 2 x 10^-8^ per base per generation and a generation time of 5 years (Matz et al. 2018). We note that the mutation rate has been measured for a different coral genus (*Acropora*) and is therefore but a rough approximation (Richards et al., 2013). This makes our estimates of absolute values of time and population sizes unreliable; however, relative differences, such as population size changes or differences in migration rates, are as accurate as the models’ parameter estimates. Additionally, for both AFS-based analyses, AFS were down-projected to 80% of the actual number of genomes within each population to reduce the noise in the higher-frequency region typical for ANGSD-derived AFS.

### Identifying loci underlying population structure

To investigate the genomic organization and association of loci underlying population differentiation, we introduce a novel method that we call “LD networks”, which is an application of weighted gene co-expression network analysis (WGCNA) to genotyping data based on the measurement of genotype correlation between pairs of SNPs. WGCNA was originally developed for analysis of gene expression (Langfelder & Horvath, 2008), to identify groups of genes that exhibit similar expression dynamics across samples (*i.e*., consistently up- or down-regulated together). The LD networks approach applies this methodology to identify groups of *loci* that demonstrate correlated changes in *genotype* across samples. In this case, rather than using the correlation of expression between genes, the adjacency matrix at the core of network construction consists of the pairwise linkage disequilibrium (LD) between loci. The characteristics of resulting “SNP modules” and associated changes in allele frequency are then evaluated with respect to sample-specific traits and metadata. In a way, this methodology is equivalent to local PCA (Li & Ralph, 2019) with single-base resolution: it identifies parts of the genome that show alternative versions of population structure, but instead of grouping genomic windows (as in the local PCA method) it groups individual SNP loci.

The analysis required five steps: (1) SNPs were first identified using the same ANGSD filters as were used to characterize population structure. Posterior genotype probabilities at each locus were estimated, recorded as the probability of each individual to carry zero, one or two derived alleles with respect to the reference genome (*i.e*., 0: homozygote ancestral, 1: heterozygote, 2: homozygote derived). This information is converted into a single value of derived alleles by multiplying each probability by its corresponding genotype and adding together. (2) Genomic coordinate information was then used to filter SNPs assumed to be physically linked. For *M. cavernosa*, a linkage block was defined as any group of SNPs in which the distance between subsequent loci was less than 1000 bp. Here, the goal is not to completely remove linked sites, but rather to disregard correlations caused by close proximity of loci, to focus on longer-range correlations that might be caused by biologically interesting factors. For *S. siderea*, in the absence of a reference assembly, the relative genomic position of RAD tags is unknown, so linked sites could only be identified as those within 36 bp of each other (*i.e*., the size of a single RAD tag). For each linkage block, the SNP with the highest minor allele frequency was retained and all others were removed from further analysis. (3) Pairwise coefficient of variation (*R^2^*) was calculated for all remaining SNPs with a minor allele frequency exceeding 0.05 using the Expectation-Maximization algorithm in ngsLD (Fox et al., 2019). The resulting matrix of correlation values is equivalent, in WGCNA terms, to an unsigned adjacency matrix based on a soft thresholding power of 2. This matrix was then used as input to the WGCNA function *TOMsimilarity*, to calculate the topological overlap matrix reflecting the sharing of the “correlation neighborhood” among loci. This and all further steps in this analysis were implemented using the WGCNA package in R (Langfelder & Horvath, 2008). (4) Following network construction, distinct groups (or “modules”) of SNPs with covarying genotypes across samples were identified using unsupervised hierarchical clustering in combination with the Dynamic Tree Cut method. (5) Lastly, association of the SNP modules with sample metadata were explored. This analysis was based on module “eigengenes”, which represent the first principal component of the genotype matrix for all SNPs included in a module. Eigengene values across samples were regressed against the samples’ metadata: assignment to a specific genetic lineage, sampling site, and age cohort. Additional analysis on module-metadata association was based on the fact that not all SNPs included in the module are equal: each SNP is assigned the “module “membership”, called “kME” in WGCNA terms, which is the correlation between the SNP genotype and the eigengene of the module. Greater kME indicates that a SNP is highly representative of the module’s behavior across samples and is likely to be among the primary responders to the factor(s) driving the correlation between SNPs within the module in the first place.

A Chi-squared test was used to determine if SNPs of each module were showed evidence of clustering throughout the genome. The proportion of total SNPs expected to occur within each scaffold of the genome assembly was calculated by simply dividing the number of SNPs within each scaffold by the total number of SNPs in the dataset. For each module, the expected number of SNPs within each scaffold was then calculated as the product of the expected proportion above and the total number of module SNPs. A comparison of the observed and expected number of module SNPs within each scaffold was used as the basis for the Chi-squared significance test. Because *S. siderea* lacks a proper genome assembly, this analysis was only performed for *M. cavernosa*.

Additionally, to explore the possible functional significance of each module, we identified SNPs that fell within 2000 bp of annotated gene boundaries and conducted a rank-based gene ontology (GO) enrichment analysis using the GO_MWU method (Wright et al., 2015; https://github.com/z0on/GO_MWU). GO_MWU is particularly powerful for WGCNA modules because it performs a two-layered test: first, a Fisher’s exact test for presence-absence of GO-annotated genes in the module, and second, a within-module Mann-Whitney U-test to determine whether the GO-annotated genes rank consistently near the top of the list of module membership values. Overall significance is assessed based on random permutations. Again, this analysis was only performed for *M. cavernosa*.

### Locus-specific F_ST_

Based on the outcome of the WGCNA analysis, we hypothesized that SNPs exhibiting highest membership to each module are in fact those that are most strongly differentiated between lineages. To evaluate this hypothesis, we estimated locus-specific *F_ST_* in two ways. First, the *realSFS* function in ANGSD was used to calculate locus-specific *F_ST_* for each pairwise comparison of the four genetic lineages in each species. *F_ST_* values from all pairwise comparisons including each lineage were averaged to produce four lineage-specific *F_ST_* estimates for every SNP. Secondly, a Bayesian *F_ST_* outlier test was implemented in BayeScan v2 (Foll & Gaggiotti, 2008). Outlier loci were identified as those in which the FDR-corrected q-value based on the posterior probability of the selection model was less than 0.05. Linear correlation was used to compare SNP module membership to *F_ST_* estimates, with the expectation that loci exhibiting the highest membership to each module will also exhibit the highest *F_ST_* values for the module-associated lineage and will be identified by Bayescan as *F_ST_* outliers.

## RESULTS

### Population structure

*M. cavernosa* and *S. siderea* exhibit remarkably similar patterns of population structure across the three reef zones. Admixture analysis and IBS-based genetic distance indicate that both species comprise four distinct lineages. Two lineages occur almost exclusively in the deep habitat and two are primarily distributed across the shallow nearshore and outer reef habitats (Figure 1B, 1C; S3). Admixed individuals with more than one source of ancestry are rare—only seven and two individuals of *M. cavernosa* and *S. siderea*, respectively, exhibit more than 25% ancestry from a secondary lineage (Figure 1B, 1C). Global *F_ST_* between lineages ranges from 0.06-0.19 in *M. cavernosa* and 0.07-0.25 in *S. siderea* (Figure 2A; Table S1). There is no consistent spatial pattern in the magnitude of differentiation—in *M. cavernosa*, the two sympatric deep populations exhibit the highest *F_ST_* of all comparisons, and in *S. siderea*, the lowest *F_ST_* is observed between one of the deep and one of the shallow lineages (Figure 2A; Table S1).

Comparing the genetic structure of adult and juvenile cohorts within each habitat reveals a degree of local adaptation. In *M. cavernosa*, juveniles of light and dark purple ancestry occur at relatively equal abundance at the nearshore and offshore sites (7:10 and 9:9, respectively; Figure 1C), while adult populations in each environment shift predominantly to one or the other. Specifically, adults of dark purple ancestry outnumber those of light purple ancestry 13:4 in the nearshore environment, but are outnumbered 5:12 in the offshore environment (Figure 1C).

**Figure 2.**
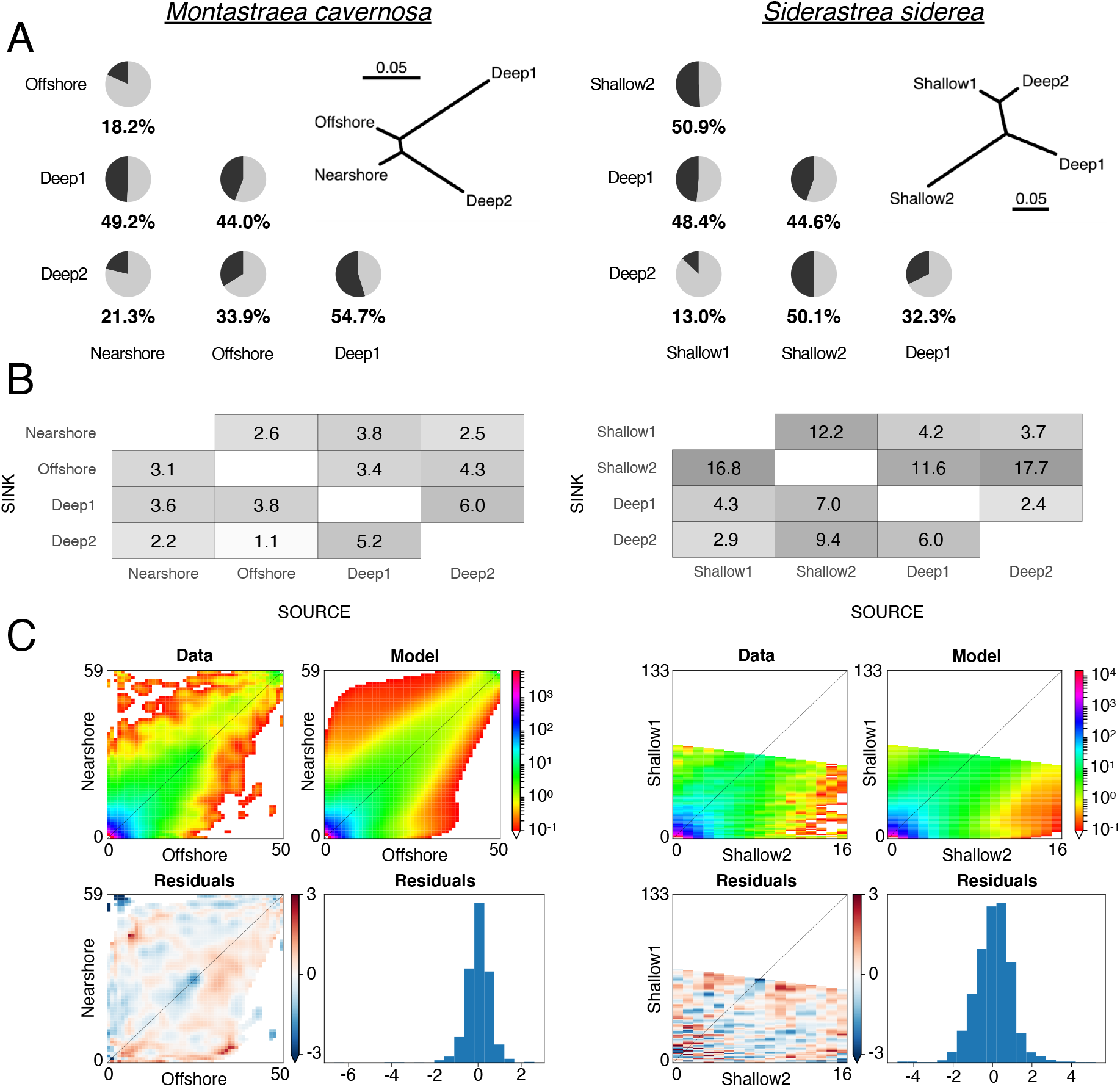
Demographic modeling indicates reduced gene flow in substantial proportion of the genome. **(A)** Pie charts depict the proportion of the genome experiencing lower introgression rate between each population pair compared to the rest of the genome (“islands of differentiation”). Distance trees reflecting the global mean pairwise *F*_ST_ between lineages is displayed in the top right corner of this panel. **(B)** Fold change by which the introgression is reduced at the islands of differentiation. Box shading reflects the magnitude of reduction. **(C)** Examples of the diagnostic plots to show goodness-of fit of the demographic models. For each species, the top two panels show the observed allele frequency spectra AFS (left) and the modeled AFS (right); the lower two panels show model errors plotted as residuals in the AFS space (left) and as a histogram (right).

Assuming a constant recruitment rate across the two generations, we can calculate the relative fitness of each ancestral lineage in the two environments as:

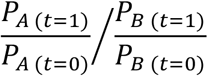

where *P_A_* is the frequency of the dominant lineage and *P_B_* is the frequency of the non-dominant lineage before (*t* = 0; juvenile) and after (*t* = 1; adult) a selection episode. Here, we find that individuals of dark purple ancestry are 2.3 times more likely to reach adulthood in the nearshore environment, while those of light purple ancestry are 2.4 times more likely to reach adulthood in the offshore environment. Additionally, only one adult colony with ancestry to either of the two deep lineages (*i.e*., light or dark green) is found in the shallow sites, and vice versa from shallow to deep. Juveniles also demonstrate nearly-perfect segregation by depth (we found only one juvenile of a shallow lineage at the deep site). To emphasize this pattern of environmental specialization and maintain clarity throughout the remainder of the text, we will refer to the four *M. cavernosa* lineages as follows: Nearshore (dark purple), Offshore (light purple), Deep1 (dark green), Deep2 (light green) (Figure 1C).

In *S. siderea*, the separation by depth is similarly pronounced among adults but not among juveniles. About half of all juveniles at depth are of dark purple ancestry, which is the most common among shallow adults and entirely dominates the shallow juvenile populations. In fact, we did not find any juveniles of light purple ancestry, the other shallow-exclusive lineage. In such a situation, there is no evidence of specialization between the two shallow lineages across the nearshore and offshore habitats, but again a notable divergence across depth. We will be referring to *S. siderea* lineages as follows throughout the remainder of the text: Shallow1 (dark purple), Shallow2 (light purple), Deep1 (dark green), Deep2 (light green) (Figure 1C).

### Demographic modeling of lineage pairs

In both species, demographic models that most accurately reproduce pairwise allele frequency spectra (AFS) between subpopulations (Figure S4, S5) all share one important feature. In every case, the model with highest likelihood includes the presence of “islands of differentiation” experiencing lower introgression rates than the rest of the genome, with the best model without this parameter ranking not higher than 20^th^ from the top by the Akaike Information Criterion. Figure 2A depicts the proportion of the genome attributable to such islands for each pairwise comparison, and Figure 2B provides the fold reduction in introgression rates within the islands compared to the rest of the genome.

In *M. cavernosa*, introgression between lineages within the islands of differentiation is reduced in 18.2-54.7 percent of the genome by a factor of 1.1-6.0. Similarly, in *S. siderea*, introgression is reduced in 13.0-50.9 percent of the genome by a factor of 2.4-17.7 (Figure 2B, 3).

**Figure 3.**
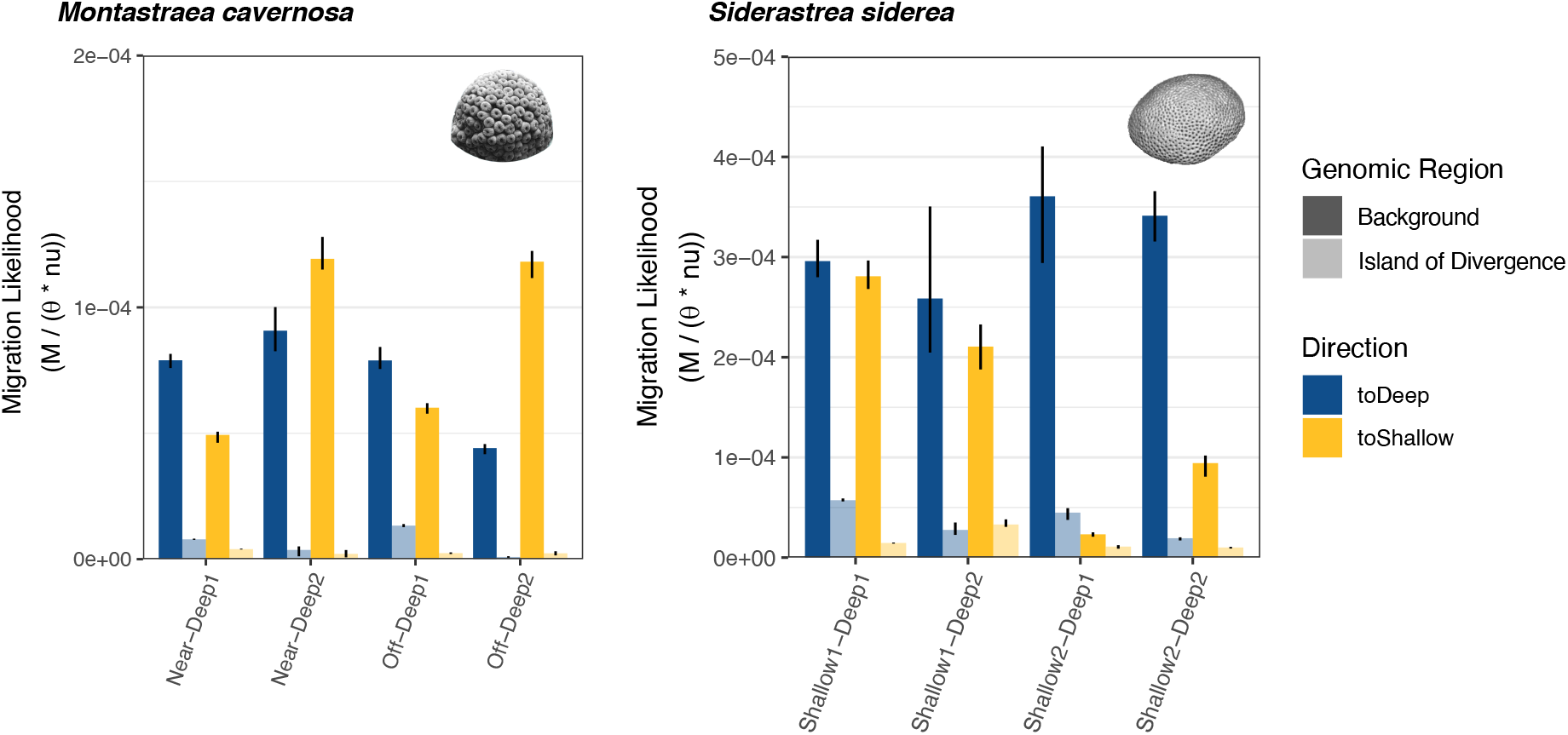
Asymmetrical introgression from shallow to deep. Median estimates of asymmetrical migration parameters between shallow and deep lineages output from bootstrapped pairwise demographic modeling for *M. cavernosa* (left) and *S. siderea* (right). Shallow-to-deep migration estimates are colored blue and deep-to-shallow estimates are colored yellow. Color saturation indicates if the estimate is associated with the background (full color) or “islands of divergence” (faded) portion of the genome. Error bars depict the lower and upper quartiles of the parameter estimates, based on all bootstrapped modeling runs.

### Effective population size changes through time

Genetic lineages of both species show broadly similar profiles of effective population size (*N*_e_) through time (Figure 4). In both species, all lineages first show *N*_e_ increase between 500 and 200 KYA. After that, three lineages in *M. cavernosa* and one lineage in *S. siderea* experience a reduction around 50-100 KYA, followed by re-expansion to about 2.5x the *N*_e_ before the reduction. Lastly, two lineages in both *M. cavernosa* and *S. siderea* show decline over the past few thousand years. The ancient and recent expansions with similar time stamps are also inferred by *Moments* models (Figure S4, S5), although these models don’t show population reductions possibly because *Moments* models could only incorporate up to three population size changes. The difference in *N*_e_ among lineages are concordant between StairwayPlot and *Moments* inference. In particular, the Shallow2 lineage of *S. siderea* has the smallest *N*_e_ both in StairwayPlot and in *Moments* and even experiences a long-term reduction according to the *Moments* models in every pairwise comparison (Figure S5). Notably, this is the least abundant lineage among our adult samples, and we did not find a single associated juvenile at any site.

**Figure 4.**
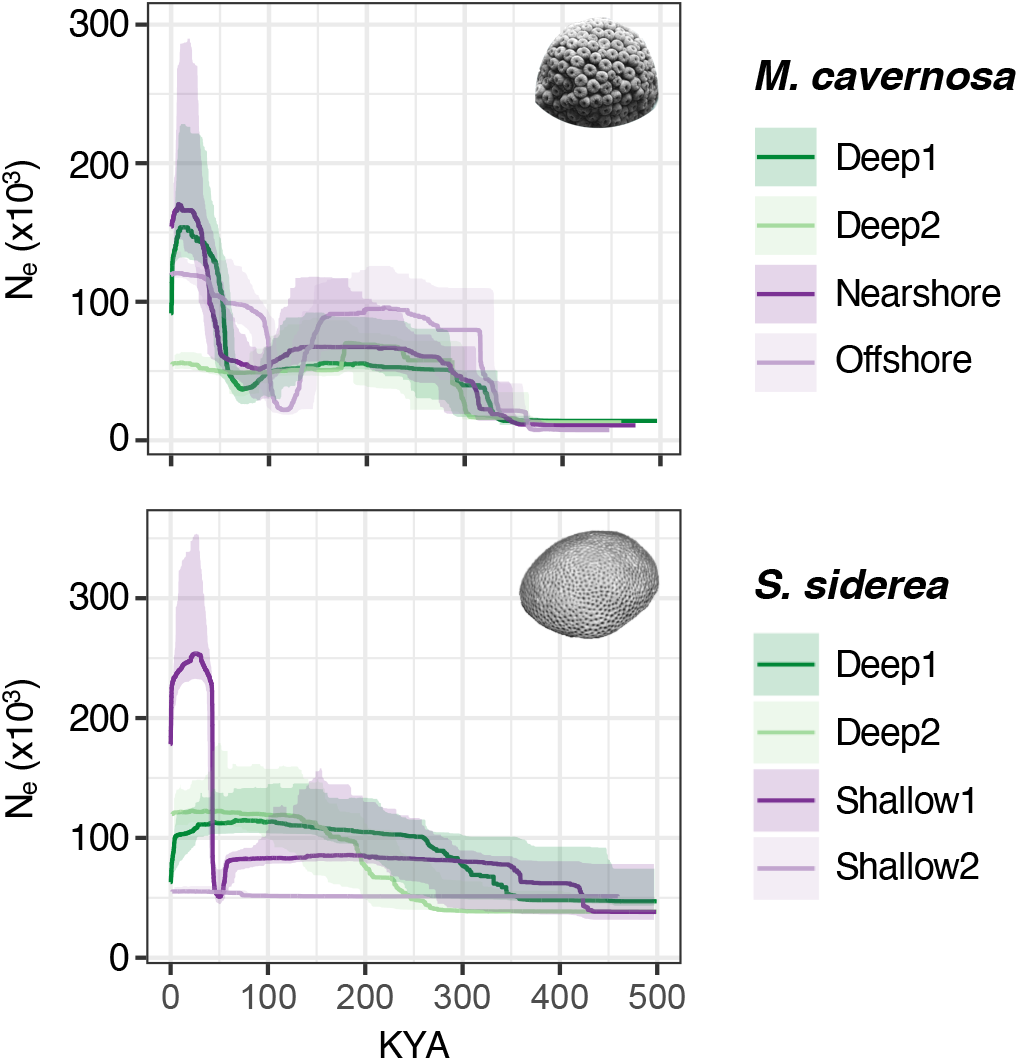
Changes in effective population size. History of effective population size changes for each of the four lineages of *M. cavernosa* (top) and *S. siderea* (bottom), as output from StairwayPlot. Colors correspond to lineage assignments in Figure 1C. 75% confidence intervals are displayed as bounding ribbons.

### Loci underlying population structure

The specific loci underlying the distinction between identified lineages were investigated using the novel LD networks methodology, which is an adaptation of the Weighted Gene Coexpression Network Analysis (WGCNA) to identify groups (“modules”) of SNPs that covary across samples. This unsupervised analysis generated exactly four distinct SNP modules for each of the two species, labeled as colors in Figure 5A. For both species, each of the four SNP module “eigengenes” (*i.e*., the weighted average genotype of module SNPs) is strongly correlated with each of the four identified lineages (Figure 5B; Figure S6).

**Figure 5.**
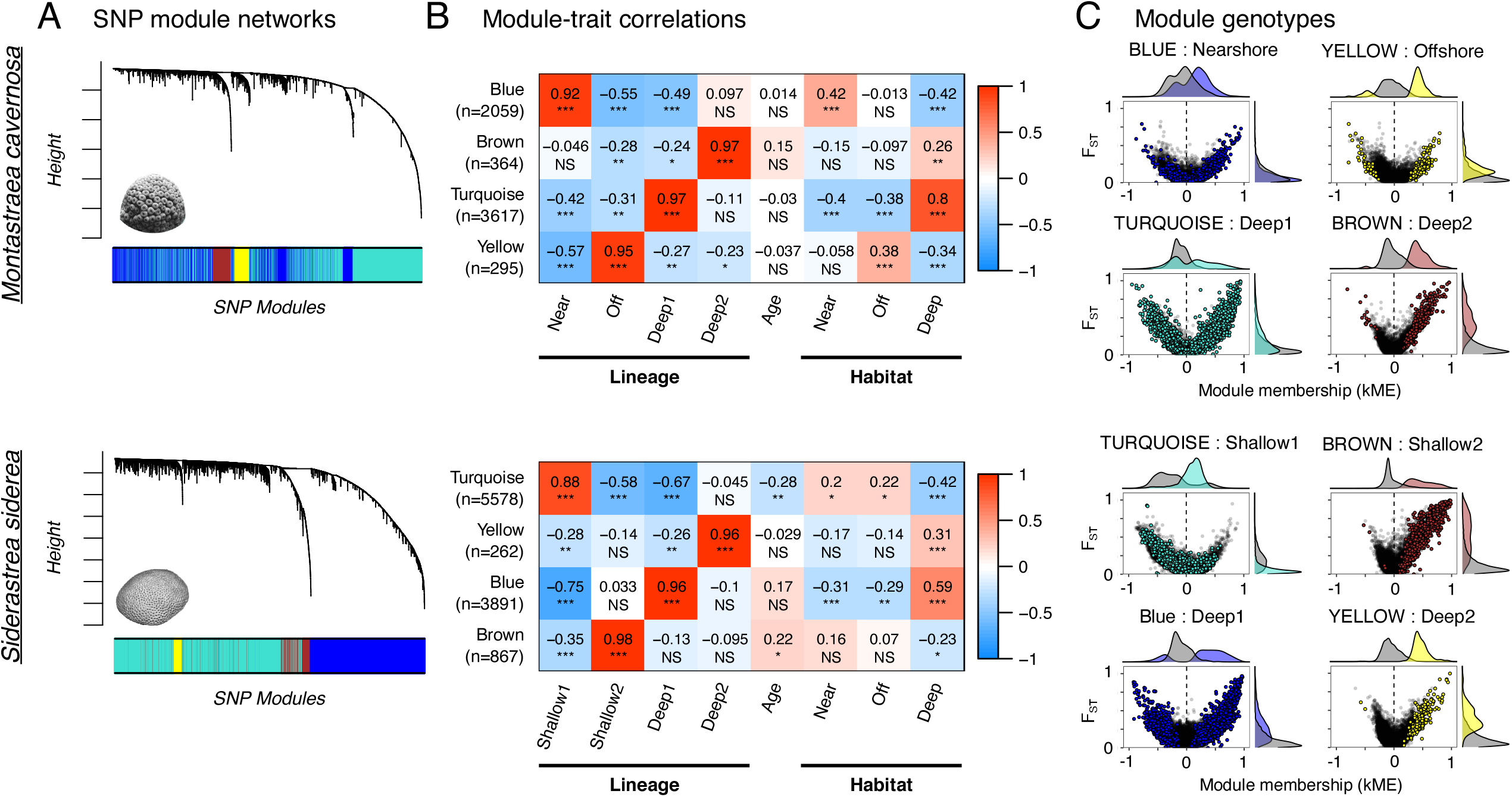
SNP modules reflect loci underlying population differentiation. **(A)** Hierarchical trees display the interconnectedness of loci based on the measure of correlation (R^2^). Correlated SNPs are parsed into distinct modules, represented below the trees as colors. **(B)** SNP module eigengenes. or the weighted average genotype of SNPs within each module, are correlated with sample traits. Correlation coefficients are provided with associated p-values in parentheses. A positive correlation (red) indicates that the majority of samples positive for the trait had higher derived allele frequency at the loci in the module, while a negative correlation (blue) indicates the opposite. Significance of module-trait correlations are reflected in the intensity of the box fill color and also under the correlation coefficient, as follows: ***: p < 0.0001. **: p < 0.001. *: p < 0.05. NS: not significant. **(C)** For the four SNP modules (four panels) of each species. SNP module membership is compared to locus-specific pairwise *F*_ST_ estimates of all lineage pairs that include the lineage most correlated with the SNP module (panel B). SNPs assigned to each module are highlighted in the color corresponding the module name. Density plots along the axes illustrate the tendency for SNPs with highest membership to each module (distance from 0 on the x-axis) to also exhibit the highest *F*_ST_ between the correlated lineage and all other lineages (y-axis).

To corroborate that the SNP modules comprise loci that underlie the differentiation of each lineage from the others, *F_ST_* was calculated for each of the SNPs included in this analysis. In both species, loci with stronger genotype correlation (either positive or negative) to each SNP module eigengene (*i.e*., stronger membership to each module) also exhibit greater *F_ST_* values (*M. cavernosa: R^2^* = 0.24, p < 0.001, *S. siderea: R^2^* = 0.30, p < 0.001; Figure 5C). As expected, the loci identified by BayeScan as putatively under selection based on an *F_ST_* outlier test fall at the extreme bounds of this association, where the absolute value of SNP-module correlation is greater than 0.5 (Figure 6).

**Figure 6.**
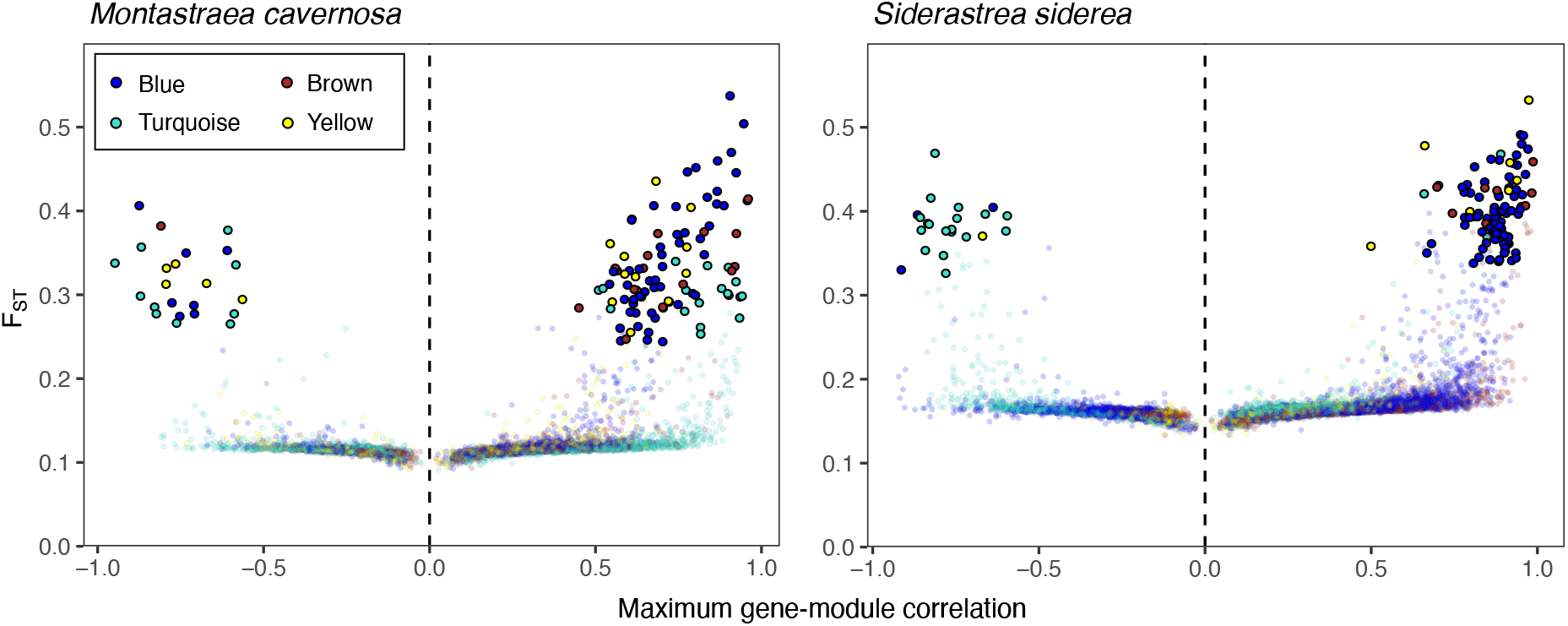
SNP module membership recapitulates *F*_ST_. Each SNP is represented as a point, colored based on the WGCNA module to which it is most strongly correlated. Correlation to this module is reflected by its position along the x-axis. Positive correlation indicates SNPs that exhibit genotypic changes across samples in the same direction as the module eigengene (*i.e*., the weighted average genotype of all SNPs in the module). Negative correlation indicates SNPs that exhibit genotypic changes in the opposite direction as the module eigengene. The locus-specific component of *F*_ST_, as calculated by BayeScan, is reflected on the y-axis. SNPs identified as putatively under selection based on an F_ST_ outlier test are bolded.

Interestingly, module-specific loci do not seem to exhibit any clustering in the genome of *M. cavernosa* (due to the lack of an annotated reference genome, this could not be assessed for *S. siderea*). Instead, these loci seem to be scattered randomly across genomic scaffolds rather than being concentrated in a particular scaffold (Blue: χ^2^ = 982.8, df = 1375, p = 1.0; Brown: χ^2^ = 1335.7, df = 1375, p = 0.77; Turquoise: χ^2^ = 652.9, df = 1375, p = 1.0; Yellow: χ^2^ = 1385.0, df = 1375, p = 0.42). We also do not find any evidence of functional enrichment among module-forming SNPs based on the GO_MWU analysis (Wright et al., 2015).

## DISCUSSION

The data presented here reveal a picture of cryptic genetic structure that follows a strikingly consistent pattern within two widespread coral species of the Florida Keys. In both *M. cavernosa* and *S. siderea*, genetically distinct lineages exist sympatrically. Both species also exhibit a pattern of environmental specialization, particularly across depth. In *M. cavernosa*, only one individual with ancestry associated with the deep site or either of the shallow sites was found in the opposing environment. In *S. siderea*, nine juvenile colonies associated with shallow ancestry were found at the deep site, but only one adult of the same background was found. No individuals of deep ancestry were found in the shallow habitats. Despite this specialization, we find evidence of ongoing introgression between all pairs of these lineages (Figure 3), as well as putative first-generation hybrids and backcrosses (Figure 1B, 1C). Notably, introgression is uneven across the genome (Figure 2A, 2B, 3): between 13% and 54% of the genome experiences several-fold lower introgression rate than the rest, ostensibly due to some form of selection. This unequal introgression across the genome among environmentally specialized lineages is the most notable feature of the genetic system described here.

### Uneven gene flow across genome

When comparing pairs of genetic lineages, demographic models consistently indicate a substantial portion of the genome exhibits introgression rates that are reduced by a factor of up to 6.0 in *M. cavernosa* and up to 17.7 in *S. siderea* relative to the rest of the genome. Using a novel LD networks approach, we find that covarying SNPs consolidate into distinct lineage-specific modules (Figure 5A, 5B; Figure S6), where SNPs that exhibit the strongest membership to their respective module also show the highest *F_ST_* between lineages (Figure 5C, 6). Notably, there is never an obvious break between high and low in *F*_ST_ values within each module (Figure 5C, 6), indicating that the distinction between two kinds of loci suggested by our demographic models is but an approximation of the continuum of introgression rates across the genome. Still, the fact that models with heterogeneous introgression fit our data significantly better than models with uniform introgression implies the existence of a mechanism maintaining (or generating) additional genetic differentiation in a substantial portion of the genome.

The mechanisms leading to such a pattern of restricted introgression can be thought to operate at either the pre- or post-zygotic stage. Mechanisms of pre-zygotic isolation can be diverse, ranging (in corals) from spawning asynchrony to gametic incompatibility (Levitan et al., 2004; Levitan & Ferrell, 2006; Ohki et al., 2015). However, all pre-zygotic mechanisms are expected to generate genome-wide divergence rather than the heterogeneous divergence across the genome.

Rather, post-zygotic factors that reduce the fitness of hybrid larvae or adults are a more likely explanation, since this type of incompatibility is expected to be restricted to certain genes rather than the whole genome. One possibility is strong spatially varying selection, preventing introgression of locally adaptive alleles across environmental boundaries (Feder et al., 2012; Rose et al., 2018). However, it is difficult to imagine how locally adaptive alleles could be so numerous (more than 50% of the genome in some cases), how can they be spread evenly across the genome, and not be associated with any specific biological functions. Moreover, this explanation clearly does not apply to the sympatric depth-specialized lineages. Another, more likely, possibility is that introgression is prevented by intra-genomic incompatibility among alleles (Dobzhansky-Muller or DM incompatibilities; Orr (1996)) such that certain combinations of alleles at different loci that arise during lineage mixing are maladaptive and therefore selected against when a first-generation hybrid starts backcrossing to one of the parental lineages. While the theoretical framework for such a mechanism was first introduced by Dobzhansky (1937) and Muller (1942) over 75 years ago, only recently has genomics research demonstrated empirical support (Powell et al., 2020; reviewed in Presgraves, 2010). The two coral species studied here represent good study systems for future research to investigate the role of DM incompatibilities in creating genetic barriers.

### Population size changes through time

Inferred *N*_e_ values and associated time stamps (Figure 4; S4, S5) must be viewed with caution because of high uncertainty in the generation time and especially the mutation rate in these corals. Still, fold-change in *N*_e_ and relative placement of expansions and declines along the time axis do not depend on these assumptions and can therefore be evaluated more reliably. It is tempting to associate declines around 50-100 KYA with glacial cycles and subsequent expansions with the stability of the last interglacial, but this is only a tentative suggestion given our uncertainty about absolute time. It is also worth mentioning that the “expansion” after 50 KYA might not truly be an increase in population size but could instead reflect the onset of introgression between lineages. Such a case would lead to higher genetic diversity and therefore higher *N*_e_ within each lineage. The most recent decline in two *M. cavernosa* and two *S. siderea* lineages may appear to reflect the precipitous decline of Florida reefs in the last 50 years (Toth et al., 2019); however, StairwayPlot is unlikely to resolve such recent events (X. Liu & Fu, 2015). More likely, this *N*_e_ decline reflects the longer-term decline of Florida Keys reefs since the flooding of the Florida Bay a few thousand years ago (Toth et al., 2018).

### Environmental specialization across depth

A substantial body of literature investigating vertical connectivity in corals has accumulated in response to renewed interest in the ‘deep reef refugia’ hypothesis. This term refers to the prospect that coral populations at depth, which are protected from the most severe impacts of climate change, may sustain or repopulate degraded shallow reefs through larval subsidy (reviewed in Bongaerts et al., 2010b). However, much of this recent research has revealed a pattern of limited population connectivity across depth. In fact, studies of *M. cavernosa* have demonstrated a strong genetic barrier between deep and shallow populations in the Florida Keys, The Bahamas, U.S. Virgin Islands and Belize (Brazeau et al., 2013; Eckert et al., 2019; Serrano et al., 2014), although not in the NW Gulf of Mexico or Bermuda (Serrano et al., 2014; Studivan & Voss, 2018). Similarly, other species both in the Caribbean and Indo-Pacific show limited dispersal between depth zones, but with variation between regions and species (Bongaerts et al., 2017; Serrano et al., 2016; van Oppen et al., 2011).

This study provides another example of genetic differentiation across depth, suggesting limited practicality for the ‘deep reef refugia’ hypothesis in the Florida Keys. Furthermore, the notable discrepancy between cross-depth migrants of the juvenile and adult life stages in *S. siderea* offers insight into the mechanism underlying this divergence over a single generation. The presence of juveniles with shallow ancestry at the deep site indicates that successful larval recruitment from shallow to deep sites occurs at an appreciable frequency. However, the lack of adults with the same shallow ancestry implies one of two possibilities. The first and perhaps more likely scenario is that the majority of juveniles are not able to survive the deep conditions and therefore do not reach adulthood, a straightforward signature of spatially varying selection. This could be due to a number of environmental factors that differ significantly across depth, but light availability in particular has been shown to strongly affect coral physiology (Lesser et al., 2010; Suggett et al., 2013; Treignier et al., 2008; Villinski, 2003) and likely plays a role in filtering maladapted genotypes from their nonnative habitats. Other selective factors may include temperature conditions, particularly in regions of frequent upwelling, or variations in viable symbiont communities, which are known to be structured significantly by depth (Bongaerts et al., 2015; Eckert et al., 2020; Lesser et al., 2010). Alternatively, because this study represents only a single snapshot of a continuous process, a second possibility is that we have captured a recent shallow-to-deep colonization event in *S. siderea*, which in time may yield a third sustainable adult population at depth.

### Environmental specialization across reef zones

In addition to the pronounced genetic barrier across depth, there is more subtle evidence of a selective gradient between nearshore and offshore reef zones. Particularly in *M. cavernosa*, comparison of adult and juvenile populations indicates that individuals assigned to the Nearshore or Offshore lineage are more than twice as likely to reach adulthood in their local habitat than the alternative lineage. This finding aligns with previous research demonstrating local adaptation across the cross-shelf gradient, a pattern that is often associated with differences in thermal tolerance. Evidence suggests that corals from nearshore reef habitats, which experience extreme daily and seasonal seawater temperature fluctuations, are more resilient to warming than their outer reef conspecifics, which experience more thermally stable conditions (Barshis et al., 2013; Castillo et al., 2012; Kenkel & Matz, 2016; Palumbi et al., 2014). In the Florida Keys, this pattern is compounded by a cross-shelf gradient in water quality, characterized by decreasing turbidity and nutrient concentrations moving away from shore (Lapointe et al., 2019; Lirman & Fong, 2007). Variation in water quality parameters, such as overall nutrient load, dissolved nutrient stoichiometry, and the concentration of suspended particulate can dramatically alter coral physiology (Allgeier et al., 2020; Anthony & Fabricius, 2000; Koop et al., 2001) and has been shown to diminish coral resistance to bleaching and disease (DeCarlo et al., 2020; Vega Thurber et al., 2014; Wiedenmann et al., 2013). It is likely that the combined pressure of these cross-shelf gradients in temperature and water quality conditions is responsible for the observed genetic specialization to each reef zone.

In *S. siderea*, while there is no evidence of a specialization across shallow nearshore and offshore reef zones, it is notable that every pairwise demographic model indicated a recent reduction in effective population size for the Shallow2 lineage (Figure S5). This pattern, coupled with the fact that no juveniles of this lineage were found, suggests that this population may be facing competitive exclusion in this region of the Florida Keys Reef Tract. By contrast, the ubiquity of the larger shallow lineage (Shallow1) across the two shallow sites and evidence of successful recruitment to the deep suggests that this lineage may be capitalizing on a selective advantage throughout this region.

### Origin of lineages

The fact that two phylogenetically divergent species (belonging to different families of the order Scleractinia) show such strikingly similar patterns of genetic subdivision (Figure 1) and demographic history (Figure 4; S4, S5) suggests that the mechanism that gave rise to their genetic structure might be linked to the properties of the environment that they share, (*i.e*., the Florida Keys Reef Tract). The Florida Keys Reef Tract is unusual among Caribbean reef systems in that it has been influenced by the tidal flow from the Florida Bay over the past few thousand years, which can be detrimental for reef development (Toth et al., 2018). Moreover, it can receive immigrant larvae from distant locations that are not directly connected to each other, such as The Bahamas and Mexico (Schill et al., 2015). It is possible that a combination of these factors led to accumulation and sympatric survival of several distinct lineages that originated in allopatry elsewhere in the Caribbean.

We expect that pre-zygotic isolation (at least for some time), local adaptation, and accumulation of DM incompatibilities all play a role in generating and sustaining genetic lineages; although, more research involving whole-genome sequencing and controlled crosses is required to confirm this conjecture. Interestingly, thus far our *Moments* models support the involvement of a period of isolation for only four of the lineage pairs in *S. siderea* (Deep1-Shallow1, Deep1-Shallow2, Deep1-Deep2, Deep2-Shallow2) and no lineage pairs in *M. cavernosa* (Figure S4, S5). Our data also indicate that new deep lineages originate from shallow rather than from other deep lineages based on the following evidence: (i) the two deep lineages in both species are the most highly divergent of all lineage pairs (Figure 2A; Table S1), (ii) in both species one of the deep lineages is closely related to one of the shallow ones, both in terms of *F*_ST_ (Figure 2A) and time since divergence (Figure S4, S5).

Notably, in *M. cavernosa* the closest lineage to one of the deep ones is the Nearshore lineage, not the Offshore lineage as could have been expected based on physical proximity of the offshore and the deep reef habitats (Figure 1A). It is possible therefore that nearshore-deep transition is facilitated by the shared adaptation to lower light conditions, due to higher turbidity nearshore and light attenuation at depth.

### Implications for reef evolution and management

Before the emergence of Stony Coral Tissue Loss Disease (SCTLD; Muller et al., 2020; Precht et al., 2016), the two coral species studied here were among the least vulnerable in the Florida Keys (Ruzicka et al., 2013; Toth et al., 2019). Even now, while large adult *M. cavernosa* and *S. siderea* are highly susceptible to SCTLD, young recruits of these species are still numerous and there is optimism for significant recovery. This is in sharp contrast to the once major reef-builders of Caribbean reefs, *Acropora palmata* and *Orbicella* sp., which have largely lost the capacity to replenish their populations through larval recruitment and in the Florida Keys are sustained exclusively by asexual reproduction through fragmentation (van Woesik et al., 2014). It is perhaps not coincidental that these two formerly foundational but now effectively ecologically extinct species are highly genetically uniform in the Florida Keys (Devlin-Durante & Baums, 2017; Manzello et al., 2019), while the two relatively successful species studied here demonstrate similar subdivision into environmentally-specialized, semi-isolated genetic lineages. Perhaps the capacity to split and specialize allows the species as a whole to accumulate broader adaptive genetic variation that fuels evolutionary rescue in times of change. This is even more likely considering that despite specialization there is still appreciable gene flow between lineages (Figure 3).

With respect to ongoing coral reef management and restoration efforts, the implication of cryptic diversification and specialization within species is important, as the provenance and genetic background of nursery-propagated corals can inform managers of the most suitable environments for outplanting. In addition, hybridization between lineages *ex situ* might serve the role of assisted gene flow (AGF; Aitken & Whitlock, 2013) by facilitating the flow of adaptive variation across lineage boundaries. Notably, unlike conventional AGF that involves crossing corals from different geographic regions (Baums et al., 2019), such “local AGF” would not be restricted by regulations prohibiting the exchange and breeding of coral genotypes across national borders.

## ACKNOWLEDGEMENTS

Funding for this work was provided by National Science Foundation award OCE-1737312, and all collections were authorized under Florida Keys National Marine Sanctuaries permit #2015-071. We wish to thank Eric Bartels and Mote’s Elizabeth Moore International Center for Coral Reef Research & Restoration for invaluable field assistance and resources. The bioinformatics analysis was accomplished using computational resources provided by the Texas Advanced Computer Center.

## AUTHOR CONTRIBUTIONS

MM conceived and designed this study. MM and GD collected tissue samples in the field, and GD performed 2bRAD library preparations. ZF assembled the *Montastraea cavernosa* reference genome with annotations contributed by YL. JR and MM conducted the bioinformatic and data analysis. JR prepared the manuscript with all authors contributing to its final form.

## DATA ACCESSIBILITY

Raw 2bRAD sequence data from this study can be accessed under the NCBI BioProject Accession PRJNA679067. The *Montastraea cavernosa* reference genome is available on the Matz Lab website (https://matzlab.weebly.com/data--code.html). Bioinformatic procedures associated with the *de novo* 2bRAD methodology (https://github.com/z0on/2bRAD_denovo), *Moments*-based demographic modeling (https://github.com/z0on/AFS-analysis-with-moments), and all other data analysis procedures used in this study (https://github.com/jprippe/AdultJuv_Depth_FL) are available in the specified GitHub repositories.

## Notes

### Competing Interest Statement

The authors have declared no competing interest.

## REFERENCES

Aitken, S. N., & Whitlock, M. C. (2013). Assisted Gene Flow to Facilitate Local Adaptation to Climate Change. Annual Review of Ecology, Evolution, and Systematics, 44(1), 367–388. https://doi.org/10.1146/annurev-ecolsys-110512-135747

Allgeier, J. E., Andskog, M. A., Hensel, E., Appaldo, R., Layman, C., & Kemp, D. W. (2020). Rewiring coral: Anthropogenic nutrients shift diverse coral–symbiont nutrient and carbon interactions toward symbiotic algal dominance. Global Change Biology, 26(10), 5588–5601. https://doi.org/10.1111/gcb.15230

Anthony, K. R. N., & Fabricius, K. E. (2000). Shifting roles of heterotrophy and autotrophy in coral energetics under varying turbidity. Journal of Experimental Marine Biology and Ecology, 252(2), 221–253. https://doi.org/10.1016/S0022-0981(00)00237-9

Aranda, M., Li, Y., Liew, Y. J., Baumgarten, S., Simakov, O., Wilson, M. C., Piel, J., Ashoor, H., Bougouffa, S., Bajic, V. B., Ryu, T., Ravasi, T., Bayer, T., Micklem, G., Kim, H., Bhak, J., LaJeunesse, T. C., & Voolstra, C. R. (2016). Genomes of coral dinoflagellate symbionts highlight evolutionary adaptations conducive to a symbiotic lifestyle. Scientific Reports, 6(1), 39734. https://doi.org/10.1038/srep39734

Barkley, H. C., Cohen, A. L., McCorkle, D. C., & Golbuu, Y. (2017). Mechanisms and thresholds for pH tolerance in Palau corals. Journal of Experimental Marine Biology and Ecology, 489, 7–14. https://doi.org/10.1016/j.jembe.2017.01.003

Barshis, D. J., Ladner, J. T., Oliver, T. A., Seneca, F. O., Traylor-Knowles, N., & Palumbi, S. R. (2013). Genomic basis for coral resilience to climate change. Proceedings of the National Academy of Sciences, 110(4), 1387–1392. https://doi.org/10.1073/pnas.1210224110

Baums, I. B., Baker, A. C., Davies, S. W., Grottoli, A. G., Kenkel, C. D., Kitchen, S. A., Kuffner, I. B., LaJeunesse, T. C., Matz, M. V., Miller, M. W., Parkinson, J. E., & Shantz, A. A. (2019). Considerations for maximizing the adaptive potential of restored coral populations in the western Atlantic. Ecological Applications, 29(8), e01978. https://doi.org/10.1002/eap.1978

Bhattacharya, D., Agrawal, S., Aranda, M., Baumgarten, S., Belcaid, M., Drake, J. L., Erwin, D., Foret, S., Gates, R. D., Gruber, D. F., & others. (2016). Comparative genomics explains the evolutionary success of reef-forming corals. ELife, 5, e13288.

Boetzer, M., & Pirovano, W. (2014). SSPACE-LongRead: Scaffolding bacterial draft genomes using long read sequence information. BMC Bioinformatics, 15(1), 211.

Bongaerts, P., Frade, P. R., Hay, K. B., Englebert, N., Latijnhouwers, K. R. W., Bak, R. P. M., Vermeij, M. J. A., & Hoegh-Guldberg, O. (2015). Deep down on a Caribbean reef: Lower mesophotic depths harbor a specialized coral-endosymbiont community. Scientific Reports, 5(1), 7652. https://doi.org/10.1038/srep07652

Bongaerts, P., Riginos, C., Ridgway, T., Sampayo, E. M., Oppen, M. J. H. van, Englebert, N., Vermeulen, F., & Hoegh-Guldberg, O. (2010a). Genetic Divergence across Habitats in the Widespread Coral Seriatopora hystrix and Its Associated Symbiodinium. PLOS ONE, 5(5), e10871. https://doi.org/10.1371/journal.pone.0010871

Bongaerts, P., Ridgway, T., Sampayo, E. M., & Hoegh-Guldberg, O. (2010b). Assessing the ‘deep reef refugia’ hypothesis: Focus on Caribbean reefs. Coral Reefs, 29(2), 309–327. https://doi.org/10.1007/s00338-009-0581-x

Bongaerts, P., Riginos, C., Brunner, R., Englebert, N., Smith, S. R., & Hoegh-Guldberg, O. (2017). Deep reefs are not universal refuges: Reseeding potential varies among coral species. Science Advances, 3(2), e1602373. https://doi.org/10.1126/sciadv.1602373

Brazeau, D. A., Lesser, M. P., & Slattery, M. (2013). Genetic Structure in the Coral, *Montastraea cavernosa*: Assessing Genetic Differentiation among and within Mesophotic Reefs. PLoS ONE, 8(5). https://doi.org/10.1371/journal.pone.0065845

Bryant, D. M., Johnson, K., DiTommaso, T., Tickle, T., Couger, M. B., Payzin-Dogru, D., Lee, T. J., Leigh, N. D., Kuo, T.-H., Davis, F. G., Bateman, J., Bryant, S., Guzikowski, A. R., Tsai, S. L., Coyne, S., Ye, W. W., Freeman, R. M., Peshkin, L., Tabin, C. J., … Whited, J. L. (2017). A Tissue-Mapped Axolotl De Novo Transcriptome Enables Identification of Limb Regeneration Factors. Cell Reports, 18(3), 762–776. https://doi.org/10.1016/j.celrep.2016.12.063

Burnham, K. P., & Anderson, D. R. (2002). Model Selection and Multimodel Inference: A Practical Information-Theoretic Approach (2nd ed.). Springer-Verlag. https://doi.org/10.1007/b97636

Campbell, M. S., Law, M., Holt, C., Stein, J. C., Moghe, G. D., Hufnagel, D. E., Lei, J., Achawanantakun, R., Jiao, D., Lawrence, C. J., Ware, D., Shiu, S.-H., Childs, K. L., Sun, Y., Jiang, N., & Yandell, M. (2014). MAKER-P: A Tool Kit for the Rapid Creation, Management, and Quality Control of Plant Genome Annotations. Plant Physiology, 164(2), 513–524. https://doi.org/10.1104/pp.113.230144

Castillo, K. D., Ries, J. B., Weiss, J. M., & Lima, F. P. (2012). Decline of forereef corals in response to recent warming linked to history of thermal exposure. Nature Climate Change, 2(10), 756–760. https://doi.org/10.1038/nclimate1577

DeCarlo, T. M., Gajdzik, L., Ellis, J., Coker, D. J., Roberts, M. B., Hammerman, N. M., Pandolfi, J. M., Monroe, A. A., & Berumen, M. L. (2020). Nutrient-supplying ocean currents modulate coral bleaching susceptibility. Science Advances, 6(34), eabc5493. https://doi.org/10.1126/sciadv.abc5493

Devlin-Durante, M. K., & Baums, I. B. (2017). Genome-wide survey of single-nucleotide polymorphisms reveals fine-scale population structure and signs of selection in the threatened Caribbean elkhorn coral, Acropora palmata. PeerJ, 5, e4077. https://doi.org/10.7717/peerj.4077

Dixon, G. B., Davies, S. W., Aglyamova, G. V., Meyer, E., Bay, L. K., & Matz, M. V. (2015). Genomic determinants of coral heat tolerance across latitudes. Science, 348(6242), 1460–1462. https://doi.org/10.1126/science.1261224

Dobzhansky, T. (1937). Genetics and the Origin of Species. Columbia University Press.

Dougan, K. (2020). A comparative genomics exploration of inter-partner metabolic signaling in the coral-algal symbiosis [PhD Dissertation]. FIU Electronic Theses and Dissertations.

Duranton, M., Allal, F., Fraïsse, C., Bierne, N., Bonhomme, F., & Gagnaire, P.-A. (2018). The origin and remolding of genomic islands of differentiation in the European sea bass. Nature Communications, 9(1), 2518. https://doi.org/10.1038/s41467-018-04963-6

Eckert, R. J., Reaume, A. M., Sturm, A. B., Studivan, M. S., & Voss, J. D. (2020). Depth Influences Symbiodiniaceae Associations Among *Montastraea cavernosa* Corals on the Belize Barrier Reef. Frontiers in Microbiology, 11. https://doi.org/10.3389/fmicb.2020.00518

Eckert, R. J., Studivan, M. S., & Voss, J. D. (2019). Populations of the coral species *Montastraea cavernosa* on the Belize Barrier Reef lack vertical connectivity. Scientific Reports, 9(1), 7200. https://doi.org/10.1038/s41598-019-43479-x

English, A. C., Richards, S., Han, Y., Wang, M., Vee, V., Qu, J., Qin, X., Muzny, D. M., Reid, J. G., Worley, K. C., & Gibbs, R. A. (2012). Mind the Gap: Upgrading Genomes with Pacific Biosciences RS Long-Read Sequencing Technology. PLOS ONE, 7(11), e47768. https://doi.org/10.1371/journal.pone.0047768

Feder, J. L., Egan, S. P., & Nosil, P. (2012). The genomics of speciation-with-gene-flow. Trends in Genetics, 28(7), 342–350. https://doi.org/10.1016/j.tig.2012.03.009

Foll, M., & Gaggiotti, O. (2008). A Genome-Scan Method to Identify Selected Loci Appropriate for Both Dominant and Codominant Markers: A Bayesian Perspective. Genetics, 180(2), 977–993. https://doi.org/10.1534/genetics.108.092221

Fox, E. A., Wright, A. E., Fumagalli, M., & Vieira, F. G. (2019). ngsLD: Evaluating linkage disequilibrium using genotype likelihoods. Bioinformatics, 35(19), 3855–3856. https://doi.org/10.1093/bioinformatics/btz200

Fu, L., Niu, B., Zhu, Z., Wu, S., & Li, W. (2012). CD-HIT: Accelerated for clustering the nextgeneration sequencing data. Bioinformatics, 28(23), 3150–3152. https://doi.org/10.1093/bioinformatics/bts565

Fuller, Z. L., Mocellin, V. J. L., Morris, L. A., Cantin, N., Shepherd, J., Sarre, L., Peng, J., Liao, Y., Pickrell, J., Andolfatto, P., Matz, M., Bay, L. K., & Przeworski, M. (2020). Population genetics of the coral *Acropora millepora*: Toward genomic prediction of bleaching. Science, 369(6501). https://doi.org/10.1126/science.aba4674

Grabherr, M. G., Haas, B. J., Yassour, M., Levin, J. Z., Thompson, D. A., Amit, I., Adiconis, X., Fan, L., Raychowdhury, R., Zeng, Q., Chen, Z., Mauceli, E., Hacohen, N., Gnirke, A., Rhind, N., di Palma, F., Birren, B. W., Nusbaum, C., Lindblad-Toh, K., … Regev, A. (2011). Trinity: Reconstructing a full-length transcriptome without a genome from RNA-Seq data. Nature Biotechnology, 29(7), 644–652. https://doi.org/10.1038/nbt.1883

Gravel, S. (n.d.). Moments: Tools for demographic inference [BitBucket Repository]. Retrieved April 4, 2020, from https://bitbucket.org/simongravel/moments/src/master/

Hagedorn, M., Page, C. A., O’Neil, K., Flores, D. M., Tichy, L., Chamberland, V. F., Lager, C., Zuchowicz, N., Lohr, K., Blackburn, H., Vardi, T., Moore, J., Moore, T., Vermeij, M. J. A., & Marhaver, K. L. (2018). Successful Demonstration of Assisted Gene Flow in the Threatened Coral *Acropora Palmata* Across Genetically-Isolated Caribbean Populations using Cryopreserved Sperm. BioRxiv, 492447. https://doi.org/10.1101/492447

Hoegh-Guldberg, O., Mumby, P. J., Hooten, A. J., Steneck, R. S., Greenfield, P., Gomez, E., Harvell, C. D., Sale, P. F., Edwards, A. J., Caldeira, K., Knowlton, N., Eakin, C. M., Iglesias-Prieto, R., Muthiga, N., Bradbury, R. H., Dubi, A., & Hatziolos, M. E. (2007). Coral Reefs Under Rapid Climate Change and Ocean Acidification. Science, 318(5857), 1737–1742. https://doi.org/10.1126/science.1152509

Hoff, K. J., & Stanke, M. (2019). Predicting Genes in Single Genomes with AUGUSTUS. Current Protocols in Bioinformatics, 65(1), e57. https://doi.org/10.1002/cpbi.57

Howells, E. J., Berkelmans, R., Oppen, M. J. H. van, Willis, B. L., & Bay, L. K. (2013). Historical thermal regimes define limits to coral acclimatization. Ecology, 94(5), 1078–1088. https://doi.org/10.1890/12-1257.1

Huang, S., Kang, M., & Xu, A. (2017). HaploMerger2: Rebuilding both haploid sub-assemblies from high-heterozygosity diploid genome assembly. Bioinformatics, 33(16), 2577–2579. https://doi.org/10.1093/bioinformatics/btx220

Huerta-Cepas, J., Forslund, K., Coelho, L. P., Szklarczyk, D., Jensen, L. J., von Mering, C., & Bork, P. (2017). Fast Genome-Wide Functional Annotation through Orthology Assignment by eggNOG-Mapper. Molecular Biology and Evolution, 34(8), 2115–2122. https://doi.org/10.1093/molbev/msx148

Hughes, T. P., Barnes, M. L., Bellwood, D. R., Cinner, J. E., Cumming, G. S., Jackson, J. B. C., Kleypas, J., van de Leemput, I. A., Lough, J. M., Morrison, T. H., Palumbi, S. R., van Nes, E. H., & Scheffer, M. (2017). Coral reefs in the Anthropocene. Nature, 546(7656), 82–90. https://doi.org/10.1038/nature22901

Jouganous, J., Long, W., Ragsdale, A. P., & Gravel, S. (2017). Inferring the Joint Demographic History of Multiple Populations: Beyond the Diffusion Approximation. Genetics, 206(3), 1549–1567. https://doi.org/10.1534/genetics.117.200493

Kenkel, C. D., & Matz, M. V. (2016). Gene expression plasticity as a mechanism of coral adaptation to a variable environment. Nature Ecology & Evolution, 1(1), 1–6. https://doi.org/10.1038/s41559-016-0014

Kenkel, C. D., Almanza, A. T., & Matz, M. V. (2015). Fine-scale environmental specialization of reef-building corals might be limiting reef recovery in the Florida Keys. Ecology, 96(12), 3197–3212. https://doi.org/10.1890/14-2297.1

Kenkel, C. D., Goodbody-Gringley, G., Caillaud, D., Davies, S. W., Bartels, E., & Matz, M. V. (2013). Evidence for a host role in thermotolerance divergence between populations of the mustard hill coral (Porites astreoides) from different reef environments. Molecular Ecology, 22(16), 4335–4348. https://doi.org/10.1111/mec.12391

Kim, S. Y., Lohmueller, K. E., Albrechtsen, A., Li, Y., Korneliussen, T., Tian, G., Grarup, N., Jiang, T., Andersen, G., Witte, D., Jorgensen, T., Hansen, T., Pedersen, O., Wang, J., & Nielsen, R. (2011). Estimation of allele frequency and association mapping using nextgeneration sequencing data. BMC Bioinformatics, 12(1), 231. https://doi.org/10.1186/1471-2105-12-231

Kitchen, S. A., Crowder, C. M., Poole, A. Z., Weis, V. M., & Meyer, E. (2015). De Novo Assembly and Characterization of Four Anthozoan (Phylum Cnidaria) Transcriptomes. G3: Genes, Genomes, Genetics, 5(11), 2441–2452. https://doi.org/10.1534/g3.115.020164

Koop, K., Booth, D., Broadbent, A., Brodie, J., Bucher, D., Capone, D., Coll, J., Dennison, W., Erdmann, M., Harrison, P., Hoegh-Guldberg, O., Hutchings, P., Jones, G. B., Larkum, A. W. D., O’Neil, J., Steven, A., Tentori, E., Ward, S., Williamson, J., & Yellowlees, D. (2001). ENCORE: The Effect of Nutrient Enrichment on Coral Reefs. Synthesis of Results and Conclusions. Marine Pollution Bulletin, 42(2), 91–120. https://doi.org/10.1016/S0025-326X(00)00181-8

Koren, S., Walenz, B. P., Berlin, K., Miller, J. R., Bergman, N. H., & Phillippy, A. M. (2017). Canu: Scalable and accurate long-read assembly via adaptive k-mer weighting and repeat separation. Genome Research, 27(5), 722–736. https://doi.org/10.1101/gr.215087.116

Korneliussen, T. S., Albrechtsen, A., & Nielsen, R. (2014). ANGSD: Analysis of Next Generation Sequencing Data. BMC Bioinformatics, 75(1), 356. https://doi.org/10.1186/s12859-014-0356-4

Ladner, J. T., & Palumbi, S. R. (2012). Extensive sympatry, cryptic diversity and introgression throughout the geographic distribution of two coral species complexes. Molecular Ecology, 21(9), 2224–2238. https://doi.org/10.1111/j.1365-294X.2012.05528.x

Langfelder, P., & Horvath, S. (2008). WGCNA: An R package for weighted correlation network analysis. BMC Bioinformatics, 9(1), 559. https://doi.org/10.1186/1471-2105-9-559

Langmead, B., & Salzberg, S. L. (2012). Fast gapped-read alignment with Bowtie 2. Nature Methods, 9(4), 357–359. https://doi.org/10.1038/nmeth.1923

Lapointe, B. E., Brewton, R. A., Herren, L. W., Porter, J. W., & Hu, C. (2019). Nitrogen enrichment, altered stoichiometry, and coral reef decline at Looe Key, Florida Keys, USA: A 3-decade study. Marine Biology, 166(8), 108. https://doi.org/10.1007/s00227-019-3538-9

Lesser, M. P. (2000). Depth-dependent photoacclimatization to solar ultraviolet radiation in the Caribbean coral *Montastraea faveolata*. Marine Ecology Progress Series, 192, 137–151. https://doi.org/10.3354/meps192137

Lesser, M. P., Slattery, M., Stat, M., Ojimi, M., Gates, R. D., & Grottoli, A. (2010). Photoacclimatization by the coral *Montastraea cavernosa* in the mesophotic zone: Light, food, and genetics. Ecology, 91(4), 990–1003. https://doi.org/10.1890/09-0313.1

Levitan, D. R., & Ferrell, D. L. (2006). Selection on Gamete Recognition Proteins Depends on Sex, Density, and Genotype Frequency. Science, 312(5771), 267–269. https://doi.org/10.1126/science.1122183

Levitan, D. R., Fukami, H., Jara, J., Kline, D., McGovern, T. M., McGhee, K. E., Swanson, C. A., & Knowlton, N. (2004). Mechanisms of Reproductive Isolation Among Sympatric Broadcast-Spawning Corals of the *Montastraea Annularis* Species Complex. Evolution, 58(2), 308–323. https://doi.org/10.1111/j.0014-3820.2004.tb01647.x

Li, H., Handsaker, B., Wysoker, A., Fennell, T., Ruan, J., Homer, N., Marth, G., Abecasis, G., Durbin, R., & 1000 Genome Project Data Processing Subgroup. (2009). The Sequence Alignment/Map format and SAMtools. Bioinformatics (Oxford, England), 25(16), 2078–2079. https://doi.org/10.1093/bioinformatics/btp352

Li, H., & Ralph, P. (2019). Local PCA Shows How the Effect of Population Structure Differs Along the Genome. Genetics, 211(1), 289–304. https://doi.org/10.1534/genetics.118.301747

Lirman, D., & Fong, P. (2007). Is proximity to land-based sources of coral stressors an appropriate measure of risk to coral reefs? An example from the Florida Reef Tract. Marine Pollution Bulletin, 54(6), 779–791. https://doi.org/10.1016/j.marpolbul.2006.12.014

Liu, H., Stephens, T. G., González-Pech, R. A., Beltran, V. H., Lapeyre, B., Bongaerts, P., Cooke, I., Aranda, M., Bourne, D. G., Forêt, S., Miller, D. J., van Oppen, M. J. H., Voolstra, C. R., Ragan, M. A., & Chan, C. X. (2018). Symbiodinium genomes reveal adaptive evolution of functions related to coral-dinoflagellate symbiosis. Communications Biology, 1(1), 1–11. https://doi.org/10.1038/s42003-018-0098-3

Liu, X., & Fu, Y.-X. (2015). Exploring population size changes using SNP frequency spectra. Nature Genetics, 47(5), 555–559. https://doi.org/10.1038/ng.3254

Manzello, D. P., Matz, M. V., Enochs, I. C., Valentino, L., Carlton, R. D., Kolodziej, G., Serrano, X., Towle, E. K., & Jankulak, M. (2019). Role of host genetics and heat-tolerant algal symbionts in sustaining populations of the endangered coral Orbicella faveolata in the Florida Keys with ocean warming. Global Change Biology, 25(3), 1016–1031. https://doi.org/10.1111/gcb.14545

Martin, M. (2011). Cutadapt removes adapter sequences from high-throughput sequencing reads. EMBnet.Journal, 17(1), 10–12. https://doi.org/10.14806/ej.17.1.200

Matz, M. V. (n.d.). 2bRAD_denovo [GitHub Repository]. Retrieved April 4, 2020, from https://github.com/z0on/2bRAD_denovo

Matz, M. V. (2018). Fantastic Beasts and How To Sequence Them: Ecological Genomics for Obscure Model Organisms. Trends in Genetics, 34(2), 121–132. https://doi.org/10.1016/j.tig.2017.11.002

Matz, M. V., Treml, E. A., Aglyamova, G. V., & Bay, L. K. (2018). Potential and limits for rapid genetic adaptation to warming in a Great Barrier Reef coral. PLoS Genetics, 14(4). https://doi.org/10.1371/journal.pgen.1007220

McBride, C. S., & Singer, M. C. (2010). Field Studies Reveal Strong Postmating Isolation between Ecologically Divergent Butterfly Populations. PLoS Biology, 8(10), e1000529. https://doi.org/10.1371/journal.pbio.1000529

Muller, E. M., Sartor, C., Alcaraz, N. I., & van Woesik, R. (2020). Spatial Epidemiology of the Stony-Coral-Tissue-Loss Disease in Florida. Frontiers in Marine Science, 7. https://doi.org/10.3389/fmars.2020.00163

Muller, H. J. (1942). Isolating mechanisms, evolution, and temperature. Biological Symposia, 6, 71–125.

Nielsen, R., Korneliussen, T., Albrechtsen, A., Li, Y., & Wang, J. (2012). SNP Calling, Genotype Calling, and Sample Allele Frequency Estimation from New-Generation Sequencing Data. PLOS ONE, 7(7), e37558. https://doi.org/10.1371/journal.pone.0037558

Ohki, S., Kowalski, R. K., Kitanobo, S., & Morita, M. (2015). Changes in spawning time led to the speciation of the broadcast spawning corals Acropora digitifera and the cryptic species Acropora sp. 1 with similar gamete recognition systems. Coral Reefs, 34(4), 1189–1198. https://doi.org/10.1007/s00338-015-1337-4

Oksanen, J., Blanchet, F. G., Friendly, M., Kindt, R., Legendre, P., McGlinn, D., Minchin, P. R., O’Hara, R. B., Simpson, G. L., Solymos, P., Stevens, M. H. H., Szoecs, E., & Wagner, H. (2019). vegan: Community Ecology Package. https://CRAN.R-project.org/package=vegan

Orr, H. A. (1996). Dobzhansky, Bateson, and the Genetics of Speciation. Genetics, 144(4), 1331–1335.

Orr, H. A., & Turelli, M. (2001). The Evolution of Postzygotic Isolation: Accumulating Dobzhansky-Muller Incompatibilities. Evolution, 55(6), 1085–1094. https://doi.org/10.1111/j.0014-3820.2001.tb00628.x

Palumbi, S. R. (2004). Marine Reserves and Ocean Neighborhoods: The Spatial Scale of Marine Populations and Their Management. Annual Review of Environment and Resources, 29(1), 31–68. https://doi.org/10.1146/annurev.energy.29.062403.102254

Palumbi, S. R., Barshis, D. J., Traylor-Knowles, N., & Bay, R. A. (2014). Mechanisms of reef coral resistance to future climate change. Science, 344(6186), 895–898. https://doi.org/10.1126/science.1251336

Powell, D. L., García-Olazábal, M., Keegan, M., Reilly, P., Du, K., Díaz-Loyo, A. P., Banerjee, S., Blakkan, D., Reich, D., Andolfatto, P., Rosenthal, G. G., Schartl, M., & Schumer, M. (2020). Natural hybridization reveals incompatible alleles that cause melanoma in swordtail fish. Science, 368(6492), 731–736. https://doi.org/10.1126/science.aba5216

Pratlong, M., Haguenauer, A., Chenesseau, S., Brener, K., Mitta, G., Toulza, E., Bonabaud, M., Rialle, S., Aurelle, D., & Pontarotti, P. (2017). Evidence for a genetic sex determination in Cnidaria, the Mediterranean red coral *(Corallium rubrum)*. Royal Society Open Science, 4(3), 160880. https://doi.org/10.1098/rsos.160880

Precht, W. F., Gintert, B. E., Robbart, M. L., Fura, R., & van Woesik, R. (2016). Unprecedented Disease-Related Coral Mortality in Southeastern Florida. Scientific Reports, 6(1), 31374. https://doi.org/10.1038/srep31374

Presgraves, D. C., Balagopalan, L., Abmayr, S. M., & Orr, H. A. (2003). Adaptive evolution drives divergence of a hybrid inviability gene between two species of Drosophila. Nature, 423(6941), 715–719. https://doi.org/10.1038/nature01679

Presgraves, D. C. (2010). The molecular evolutionary basis of species formation. Nature Reviews Genetics, 11(3), 175–180. https://doi.org/10.1038/nrg2718

Price, A. L., Jones, N. C., & Pevzner, P. A. (2005). De novo identification of repeat families in large genomes. Bioinformatics, 21, i351–i358. https://doi.org/10.1093/bioinformatics/bti1018

Quigley, K. M., Bay, L. K., & Oppen, M. J. H. van. (2019). The active spread of adaptive variation for reef resilience. Ecology and Evolution, 9(19), 11122–11135. https://doi.org/10.1002/ece3.5616

R Core Team. (2019). R: A Language and Environment for Statistical Computing. R Foundation for Statistical Computing. https://www.R-project.org/

Richards, Z. T., Miller, D. J., & Wallace, C. C. (2013). Molecular phylogenetics of geographically restricted Acropora species: Implications for threatened species conservation. Molecular Phylogenetics and Evolution, 69(3), 837–851. https://doi.org/10.1016/j.ympev.2013.06.020

Richards, Z. T., Berry, O., & van Oppen, M. J. H. (2016). Cryptic genetic divergence within threatened species of Acropora coral from the Indian and Pacific Oceans. Conservation Genetics, 17(3), 577–591. https://doi.org/10.1007/s10592-015-0807-0

Ronce, O., & Kirkpatrick, M. (2001). When Sources Become Sinks: Migrational Meltdown in Heterogeneous Habitats. Evolution, 55(8), 1520–1531. https://doi.org/10.1111/j.0014-3820.2001.tb00672.x

Rose, N. H., Bay, R. A., Morikawa, M. K., & Palumbi, S. R. (2018). Polygenic evolution drives species divergence and climate adaptation in corals. Evolution, 72(1), 82–94. https://doi.org/10.1111/evo.13385

Rosser, N. L. (2015). Asynchronous spawning in sympatric populations of a hard coral reveals cryptic species and ancient genetic lineages. Molecular Ecology, 24(19), 5006–5019. https://doi.org/10.1111/mec.13372

Rundle, H. D. (2002). A Test of Ecologically Dependent Postmating Isolation Between Sympatric Sticklebacks. Evolution, 56(2), 322–329. https://doi.org/10.1111/j.0014-3820.2002.tb01342.x

Ruzicka, R. R., Colella, M. A., Porter, J. W., Morrison, J. M., Kidney, J. A., Brinkhuis, V., Lunz, K. S., Macaulay, K. A., Bartlett, L. A., Meyers, M. K., & Colee, J. (2013). Temporal changes in benthic assemblages on Florida Keys reefs 11 years after the 1997/1998 El Niño. Marine Ecology Progress Series, 489, 125–141. https://doi.org/10.3354/meps10427

Schill, S. R., Raber, G. T., Roberts, J. J., Treml, E. A., Brenner, J., & Halpin, P. N. (2015). No Reef Is an Island: Integrating Coral Reef Connectivity Data into the Design of Regional-Scale Marine Protected Area Networks. PLOS ONE, 10(12), e0144199. https://doi.org/10.1371/journal.pone.0144199

Schmidt-Roach, S., Lundgren, P., Miller, K. J., Gerlach, G., Noreen, A. M. E., & Andreakis, N. (2013). Assessing hidden species diversity in the coral Pocillopora damicornis from Eastern Australia. Coral Reefs, 32(1), 161–172. https://doi.org/10.1007/s00338-012-0959-z

Serrano, X., Baums, I. B., O’Reilly, K., Smith, T. B., Jones, R. J., Shearer, T. L., Nunes, F. L. D., & Baker, A. C. (2014). Geographic differences in vertical connectivity in the Caribbean coral *Montastraea cavernosa* despite high levels of horizontal connectivity at shallow depths. Molecular Ecology, 23(17), 4226–4240. https://doi.org/10.1111/mec.12861

Serrano, X., Baums, I. B., Smith, T. B., Jones, R. J., Shearer, T. L., & Baker, A. C. (2016). Long distance dispersal and vertical gene flow in the Caribbean brooding coral *Porites astreoides*. Scientific Reports, 6(1), 21619. https://doi.org/10.1038/srep21619

Shoguchi, E., Shinzato, C., Kawashima, T., Gyoja, F., Mungpakdee, S., Koyanagi, R., Takeuchi, T., Hisata, K., Tanaka, M., Fujiwara, M., Hamada, M., Seidi, A., Fujie, M., Usami, T., Goto, H., Yamasaki, S., Arakaki, N., Suzuki, Y., Sugano, S., … Satoh, N. (2013). Draft Assembly of the *Symbiodinium minutum* Nuclear Genome Reveals Dinoflagellate Gene Structure. Current Biology, 23(15), 1399–1408. https://doi.org/10.1016/j.cub.2013.05.062

Simão, F. A., Waterhouse, R. M., Ioannidis, P., Kriventseva, E. V., & Zdobnov, E. M. (2015). BUSCO: Assessing genome assembly and annotation completeness with single-copy orthologs. Bioinformatics, 31(19), 3210–3212. https://doi.org/10.1093/bioinformatics/btv351

Skotte, L., Korneliussen, T. S., & Albrechtsen, A. (2013). Estimating individual admixture proportions from next generation sequencing data. Genetics, 195(3), 693–702. https://doi.org/10.1534/genetics.113.154138

Studivan, M. S., & Voss, J. D. (2018). Population connectivity among shallow and mesophotic *Montastraea cavernosa* corals in the Gulf of Mexico identifies potential for refugia. Coral Reefs, 37(4), 1183–1196. https://doi.org/10.1007/s00338-018-1733-7

Suggett, D. J., Dong, L. F., Lawson, T., Lawrenz, E., Torres, L., & Smith, D. J. (2013). Light availability determines susceptibility of reef building corals to ocean acidification. Coral Reefs, 32(2), 327–337. https://doi.org/10.1007/s00338-012-0996-7

Toth, L. T., Kuffner, I. B., Stathakopoulos, A., & Shinn, E. A. (2018). A 3,000-year lag between the geological and ecological shutdown of Florida’s coral reefs. Global Change Biology, 24(11), 5471–5483. https://doi.org/10.1111/gcb.14389

Toth, L. T., Stathakopoulos, A., Kuffner, I. B., Ruzicka, R. R., Colella, M. A., & Shinn, E. A. (2019). The unprecedented loss of Florida’s reef-building corals and the emergence of a novel coral-reef assemblage. Ecology, 100(9), e02781. https://doi.org/10.1002/ecy.2781

Trapnell, C., Roberts, A., Goff, L., Pertea, G., Kim, D., Kelley, D. R., Pimentel, H., Salzberg, S. L., Rinn, J. L., & Pachter, L. (2012). Differential gene and transcript expression analysis of RNA-seq experiments with TopHat and Cufflinks. Nature Protocols, 7(3), 562–578. https://doi.org/10.1038/nprot.2012.016

Treignier, C., Grover, R., Ferrier-Pagés, C., & Tolosa, I. (2008). Effect of light and feeding on the fatty acid and sterol composition of zooxanthellae and host tissue isolated from the scleractinian coral Turbinaria reniformis. Limnology and Oceanography, 53(6), 2702–2710. https://doi.org/10.4319/lo.2008.53.6.2702

van Oppen, M. J. H., Bongaerts, P., Underwood, J. N., Peplow, L. M., & Cooper, T. F. (2011). The role of deep reefs in shallow reef recovery: An assessment of vertical connectivity in a brooding coral from west and east Australia. Molecular Ecology, 20(8), 1647–1660. https://doi.org/10.1111/j.1365-294X.2011.05050.x

van Oppen, M. J. H., Gates, R. D., Blackall, L. L., Cantin, N., Chakravarti, L. J., Chan, W. Y., Cormick, C., Crean, A., Damjanovic, K., Epstein, H., Harrison, P. L., Jones, T. A., Miller, M., Pears, R. J., Peplow, L. M., Raftos, D. A., Schaffelke, B., Stewart, K., Torda, G., … Putnam, H. M. (2017). Shifting paradigms in restoration of the world’s coral reefs. Global Change Biology, 23(9), 3437–3448. https://doi.org/10.1111/gcb.13647

van Oppen, M. J. H., Oliver, J. K., Putnam, H. M., & Gates, R. D. (2015). Building coral reef resilience through assisted evolution. Proceedings of the National Academy of Sciences, 112(8), 2307–2313. https://doi.org/10.1073/pnas.1422301112

van Woesik, R., Scott, W. J., & Aronson, R. B. (2014). Lost opportunities: Coral recruitment does not translate to reef recovery in the Florida Keys. Marine Pollution Bulletin, 88(1), 110–117. https://doi.org/10.1016/j.marpolbul.2014.09.017

Vega Thurber, R. L., Burkepile, D. E., Fuchs, C., Shantz, A. A., McMinds, R., & Zaneveld, J. R. (2014). Chronic nutrient enrichment increases prevalence and severity of coral disease and bleaching. Global Change Biology, 20(2), 544–554. https://doi.org/10.1111/gcb.12450

Villinski, J. T. (2003). Depth-independent reproductive characteristics for the Caribbean reefbuilding coral *Montastraea faveolata*. Marine Biology, 142(6), 1043–1053. https://doi.org/10.1007/s00227-002-0997-0

Vollmer, S. V., & Palumbi, S. R. (2002). Hybridization and the Evolution of Reef Coral Diversity. Science, 296(5575), 2023–2025. https://doi.org/10.1126/science.1069524

Walker, B. J., Abeel, T., Shea, T., Priest, M., Abouelliel, A., Sakthikumar, S., Cuomo, C. A., Zeng, Q., Wortman, J., Young, S. K., & Earl, A. M. (2014). Pilon: An Integrated Tool for Comprehensive Microbial Variant Detection and Genome Assembly Improvement. PLOS ONE, 9(11), e112963. https://doi.org/10.1371/journal.pone.0112963

Wang, S., Meyer, E., McKay, J. K., & Matz, M. V. (2012). 2b-RAD: A simple and flexible method for genome-wide genotyping. Nature Methods, 9(8), 808–810. https://doi.org/10.1038/nmeth.2023

Warner, P. A., Oppen, M. J. H. van, & Willis, B. L. (2015). Unexpected cryptic species diversity in the widespread coral Seriatopora hystrix masks spatial-genetic patterns of connectivity. Molecular Ecology, 24(12), 2993–3008. https://doi.org/10.1111/mec.13225

Warren, R. L., Yang, C., Vandervalk, B. P., Behsaz, B., Lagman, A., Jones, S. J. M., & Birol, I. (2015). LINKS: Scalable, alignment-free scaffolding of draft genomes with long reads. GigaScience, 4(1). https://doi.org/10.1186/s13742-015-0076-3

Waterhouse, R. M., Seppey, M., Simão, F. A., Manni, M., Ioannidis, P., Klioutchnikov, G., Kriventseva, E. V., & Zdobnov, E. M. (2018). BUSCO applications from quality assessments to gene prediction and phylogenomics. Molecular Biology and Evolution, 35(3), 543–548.

Wiedenmann, J., D’Angelo, C., Smith, E. G., Hunt, A. N., Legiret, F.-E., Postle, A. D., & Achterberg, E. P. (2013). Nutrient enrichment can increase the susceptibility of reef corals to bleaching. Nature Climate Change, 3(2), 160–164. https://doi.org/10.1038/nclimate1661

Willis, B. L., Babcock, R. C., Harrison, P. L., & Wallace, C. C. (1997). Experimental hybridization and breeding incompatibilities within the mating systems of mass spawning reef corals. Coral Reefs, 16(0), S53–S65. https://doi.org/10.1007/s003380050242

Willis, B. L., van Oppen, M. J. H., Miller, D. J., Vollmer, S. V., & Ayre, D. J. (2006). The Role of Hybridization in the Evolution of Reef Corals. Annual Review of Ecology, Evolution, and Systematics, 37(1), 489–517. https://doi.org/10.1146/annurev.ecolsys.37.091305.110136

Wright, R. M., Aglyamova, G. V., Meyer, E., & Matz, M. V. (2015). Gene expression associated with white syndromes in a reef building coral, *Acropora hyacinthus*. BMC Genomics, 16(1), 371. https://doi.org/10.1186/s12864-015-1540-2

Yeo, S., Coombe, L., Warren, R. L., Chu, J., & Birol, I. (2018). ARCS: scaffolding genome drafts with linked reads. Bioinformatics, 34(5), 725–731.

