## Supplemental Information for "Environmental specialization and cryptic genetic divergence in two massive coral species from the Florida Keys Reef Tract"

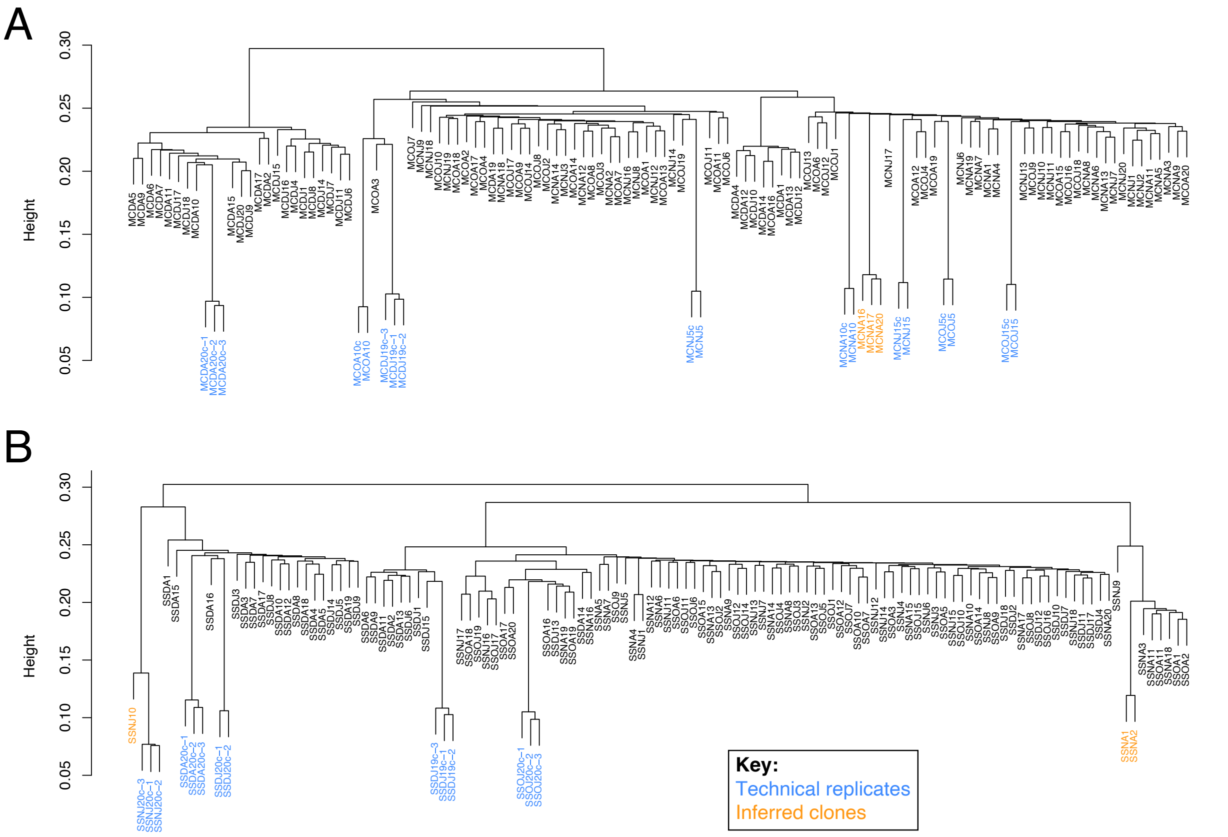


**Supplemental Figure S1. Clone detection based on identity-by-state.** Hierarchical clustering tree reflecting genetic dissimilarity based on pairwise identity-by-state (IBS) distance. Technical replicates and inferred clones are highlighted in blue and orange, respectively. Any samples exhibiting a genetic dissimilarity as low as known technical replicates were assumed to be clonemates, and only the individual of each clonal group with the greatest total read count was chosen to be retained for further analyses.

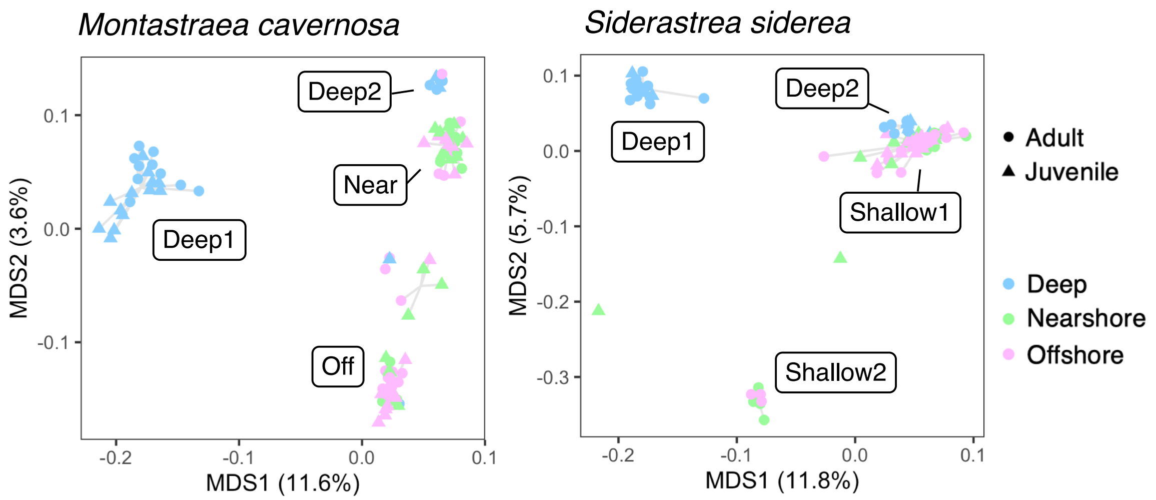


**Supplemental Figure S2. Population structure (AFS filters).** Principal coordinate analyses (PCoA) demonstrating that differentiation between the four lineages is robust to the modified SNP filters used in demographic analyses. The colors of individual points correspond to the habitats specified in Figure 1A.

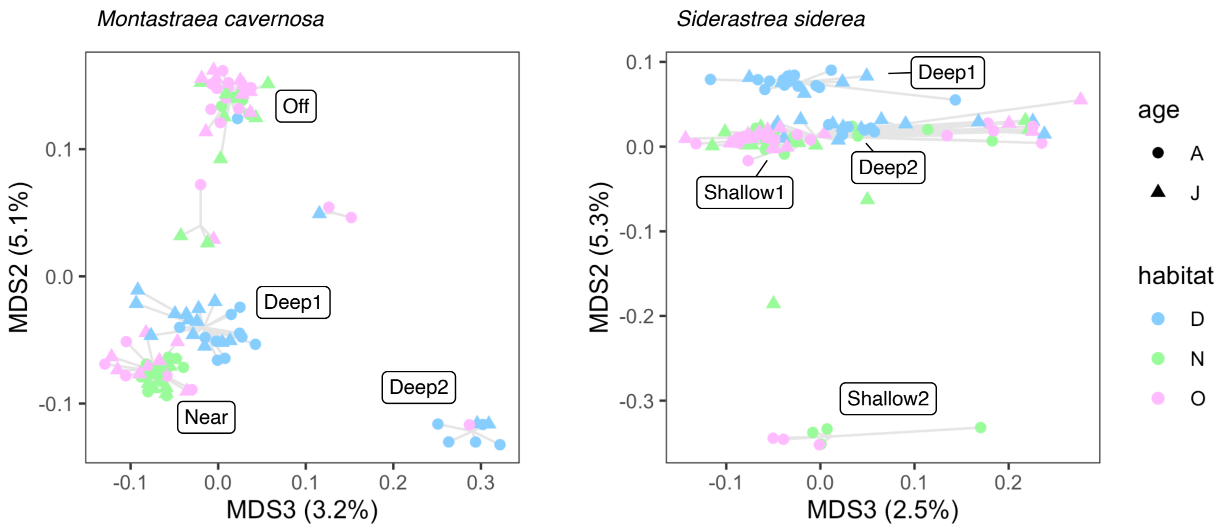


**Supplemental Figure S3. Population structure (rotated PCoA axes).** Principal coordinate analyses (PCoA) demonstrating the four distinct lineages within each species with respect to MDS3 (x-axis) and MDS2 (y-axis). The colors of individual points correspond to the habitats specified in Figure 1A.

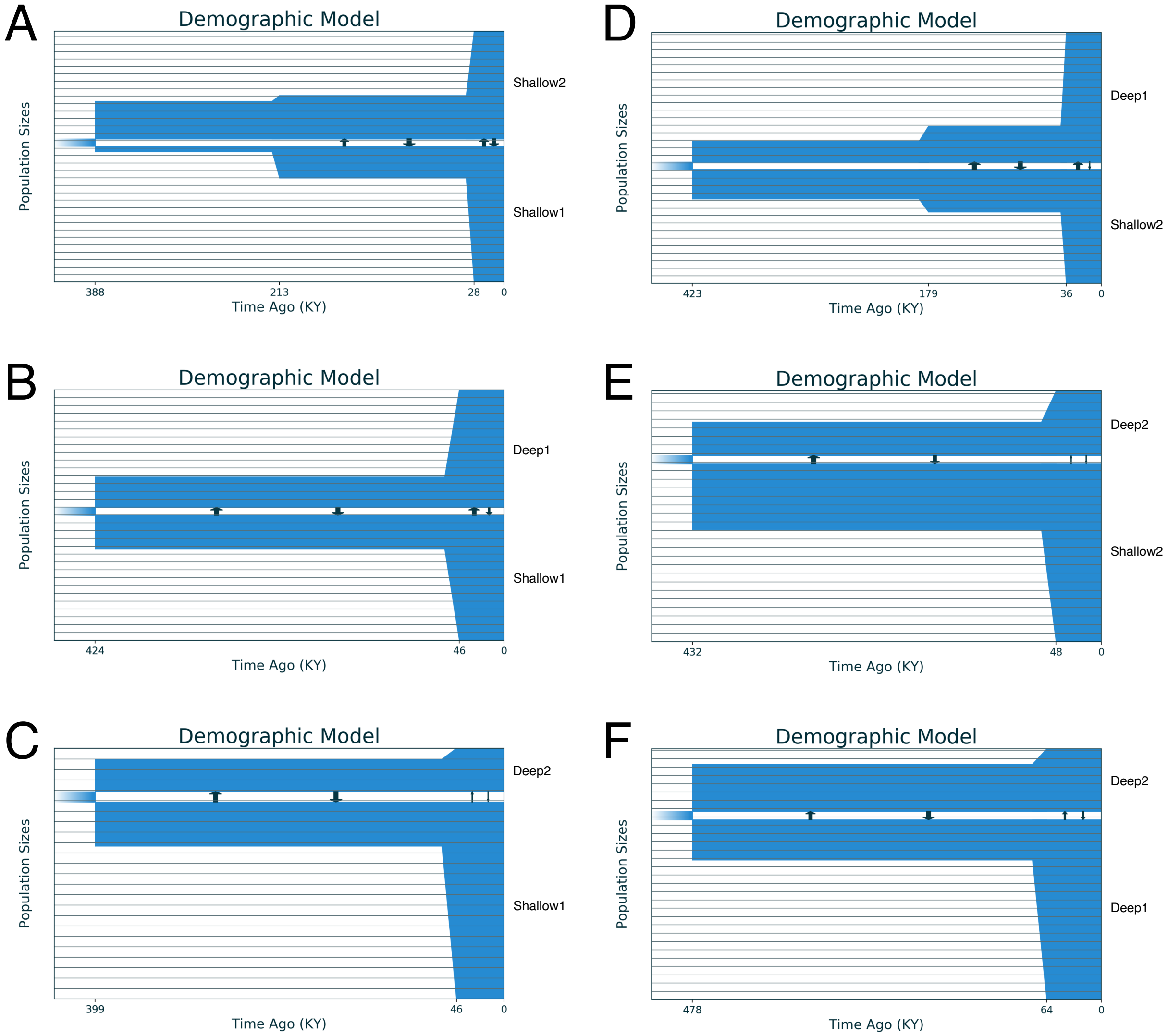


**Supplemental Figure S4. Pairwise demographic histories – *Montastraea cavernosa*.** Best-fit demographic histories based on the bootstrapped Moments modeling procedure, depicting simulated changes in effective population size and migration rates for all pairwise combinations of the four lineages in *M. cavernosa*.

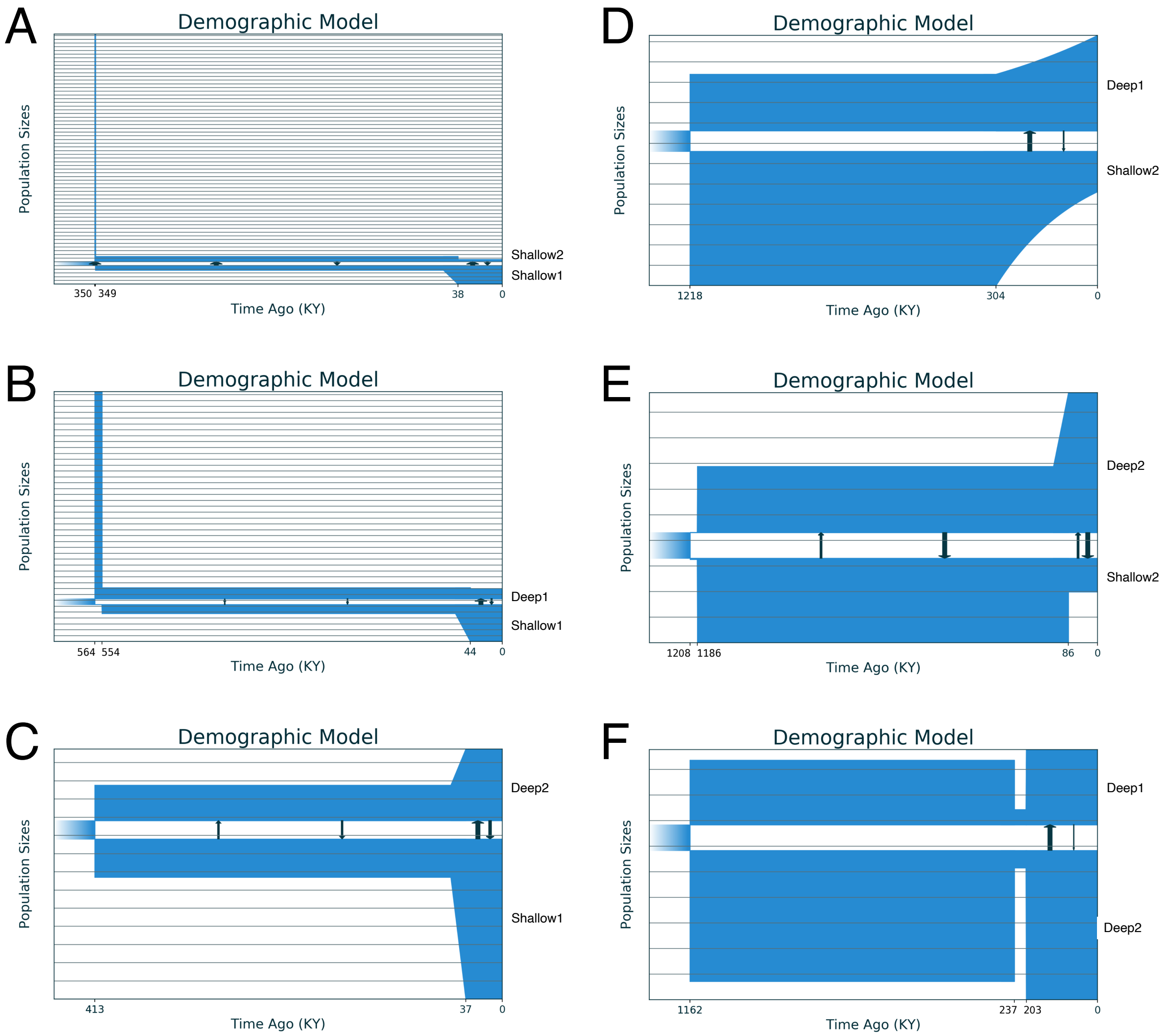


**Supplemental Figure S5. Pairwise demographic histories – *Siderastrea siderea*.** Best-fit demographic histories based on the bootstrapped Moments modeling procedure, depicting simulated changes in effective population size and migration rates for all pairwise combinations of the four lineages in *S. siderea*.

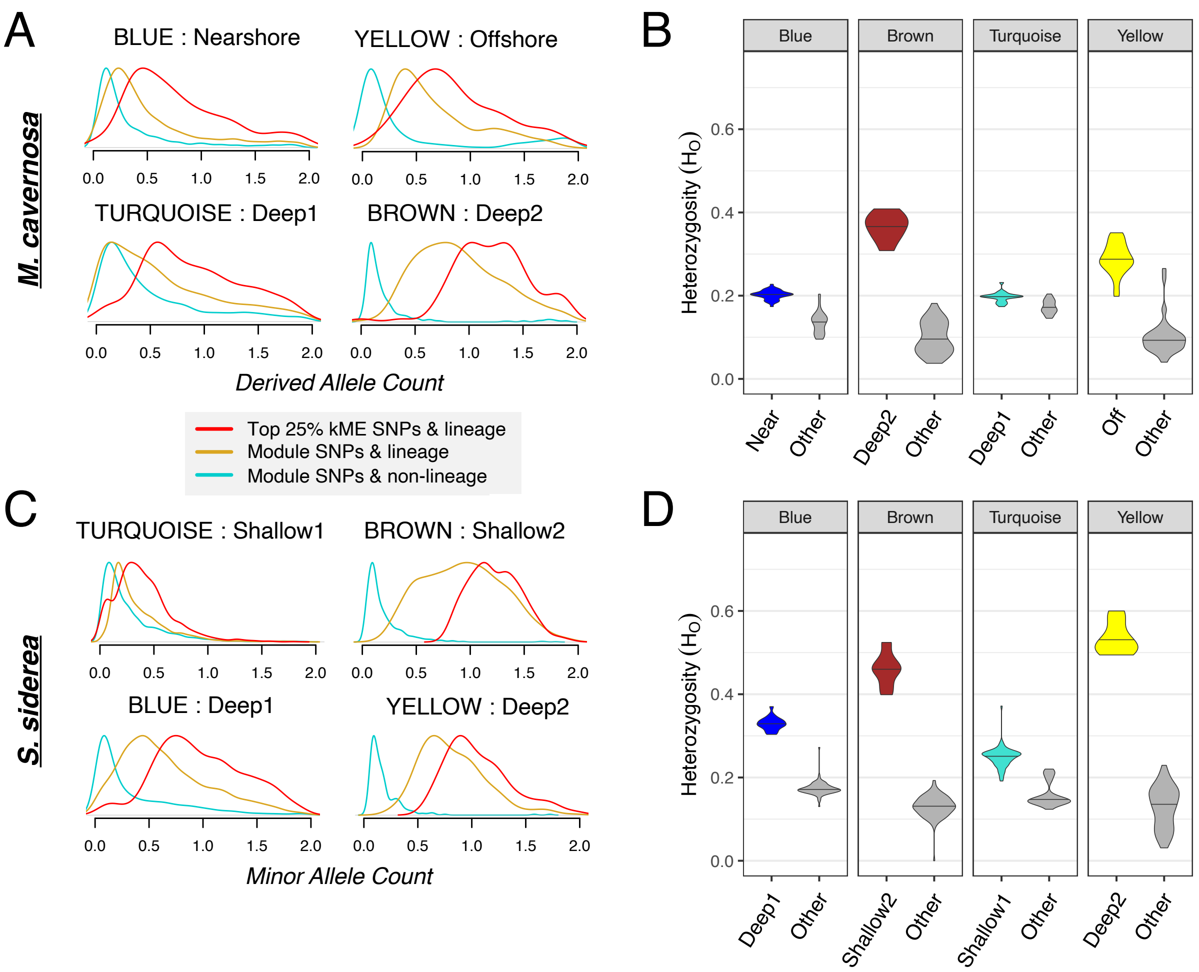


**Supplemental Figure S6. Genotypic shifts associated with SNP modules. (A and C)** For each of the four lineages within each species, density plots depict the derived/minor allele count of corresponding SNP modules that are most strongly associated with lineage membership. Gold and teal lines represent the genotypes of individuals associated and not associated with each specified lineage, respectively, at module loci. Red lines depict the genotypes of individuals associated with each lineage, but only with respect to loci with module membership in the 75$^{th}$ percentile of all module SNPs. **(B and D)** Violin plots display the observed heterozygosity of SNPs within each module (four panels) for individuals assigned to the corresponding lineage (colored) as compared to all other individuals (gray).

***M. cavernosa***

| | Nearshore | Offshore | Deep1 | Deep2 |
|---|---|---|---|---|
| Nearshore | - | | | |
| Offshore | 0.05684 | - | | |
| Deep1 | 0.160143 | 0.133304 | - | |
| Deep2 | 0.099428 | 0.124154 | 0.192365 | - |

***S. siderea***

| | Shallow1 | Shallow2 | Deep1 | Deep2 |
|---|---|---|---|---|
| Shallow1 | - | | | |
| Shallow2 | 0.230957 | - | | |
| Deep1 | 0.176799 | 0.223152 | - | |
| Deep2 | 0.067963 | 0.247109 | 0.166883 | - |

**Supplemental Table S1.** Global $F_{ST}$ estimates between all pairwise combinations of the four lineages of *M. cavernosa* and *S. siderea*.
